# Hypothermic Conditions Impair GnRH Pulse Generator Activity and Gametogenesis

**DOI:** 10.64898/2026.08.27.747452

**Authors:** Mitsue Hagihara, Daisuke Suzuki, Junko Hara, Takaya Abe, Takeshi Sakurai, Kazunari Miyamichi, Teppei Goto

## Abstract

Mammalian reproductive function is driven by arcuate kisspeptin neurons, pacemakers of gonadotropin secretion. During energy shortages, animals reallocate resources from reproduction to survival; however, the underlying neural mechanisms remain elusive. Here we used fiber photometry to chronically monitor synchronized episodes of arcuate kisspeptin neuron activity (SEs^kiss^) in adult mice under various energy-saving conditions. In both sexes, SEs^kiss^ frequency was markedly suppressed during fasting-induced torpor and pharmacologically induced hypothermia, whereas hypometabolism alone had no discernible effect. A Q neuron-induced hypothermic state (QIH) robustly suppressed SEs^kiss^, leading to impaired gamete maturation, whereas warming the body temperature during QIH fully restored SEs^kiss^ frequency. These findings demonstrate that hypothermia, rather than hypometabolism, is the primary driver of suppression of the hypothalamic reproductive axis during energy-saving conditions. This study provides insights into how thermal signals act as critical gatekeepers in the mammalian reproductive system.

## Introduction

Reproduction is highly energy-demanding; thus, there are trade-offs in energy allocation between reproduction and other metabolic functions, such as growth (*1*). In various mammalian species, including primates, a negative energy balance, such as that induced by malnutrition, suppresses reproduction. These adaptive changes are thought to arise from alterations in specific hypothalamic functions, leading to reduced activity of the hypothalamic-pituitary-gonadal (HPG) axis (*2*). Recent rodent studies have shown that the onset of female puberty is tightly regulated by food availability and the energy state of the body, which is sensed through both hormonal and neuronal pathways that modulate the hypothalamic reproductive centers (*3–5*). Similarly, delayed sexual maturation has been reported in girls undergoing intense physical training and reduced body fat (*6*, *7*). In contrast to these peripubertal contexts, it remains unclear to what extent food availability and negative energy balance affect the acute activity of the HPG axis in adulthood in both males and females. This represents a critical knowledge gap, given that extreme dieting and intermittent fasting have become increasingly popular worldwide, with a growing emphasis on health and beauty.

In mammalian reproduction, the coordinated endocrine HPG axis is primarily controlled by hypothalamic kisspeptin neurons (*8–10*). Gametogenesis is regulated by the pulsatile secretion of hypothalamic gonadotropin-releasing hormone (GnRH) and the subsequent release of pituitary gonadotropins. In turn, gonadal steroid hormones provide feedback to the hypothalamus. Accumulating evidence has shown that hypothalamic arcuate kisspeptin (ARC^kiss^) neurons, which are targets of steroid hormone-mediated negative feedback (*11–13*), play a key role in GnRH pulse generation (*14*, *15*). Previous studies have established electrophysiological recordings of multiunit activity from putative ARC^kiss^ neurons in goats (*16*, *17*) as well as cell-type-specific chronic Ca^2+^ imaging of ARC^kiss^ neural activity in freely moving mice using fiber photometry (*18*, *19*). ARC^kiss^ neurons display pulsatile activity termed <u>s</u>ynchronous <u>e</u>pisode<u>s</u> of elevated Ca^2+^ (referred to as SEs^kiss^ for simplicity (*18*)). SEs^kiss^ events are closely associated with gonadotropin secretion (*18*) and are negatively regulated by sex steroid hormones (*20*, *21*). These findings raise the question of how the temporal dynamics of SEs^kiss^ are altered under conditions of energy depletion.

How does energy deficiency lead to reproductive suppression at the level of neural circuitry? One important upstream regulator of ARC^kiss^ neurons appears to be AgRP neurons in the arcuate nucleus (ARC^Agrp^), which are orexigenic inhibitory neurons that represent a hunger state (*22*) and regulate feeding behaviors (*23*). Chemogenetic activation of ARC^Agrp^ neurons has been reported to delay the estrous cycle through the ARC^Agrp^-ARC^kiss^ neural circuit (*24*). In the context of puberty onset, our recent study combined chronic fiber photometry of ARC^kiss^ neurons with viral genetic tools to show that ARC^Agrp^ neurons reduce the frequency of SEs^kiss^ under negative energy balance (*5*). However, it remains unclear whether ARC^Agrp^ neurons exert suppressive effects on SEs^kiss^ under energy-deficient conditions in adults. More generally, the influence of metabolic challenges on the temporal dynamics of SEs^kiss^ remains unclear.

The energy state is critical not only for reproduction but also for individual survival. Various mammalian species have evolved mechanisms such as controllable hypothermia, torpor, and hibernation to reduce thermogenesis and basal metabolism, thereby enabling their survival under harsh conditions of food scarcity (*25–27*). Laboratory mice exhibit a relatively shallow form of hypothermia, called daily torpor, which is characterized by repeated bouts of reduced core body temperature (Tc) and metabolic rate (*28*, *29*). A recent study in mice demonstrated that artificial activation of pyroglutamylated RFamide peptide (Qrfp) neurons in the preoptic area can induce a state known as the Q-neuron-induced hypothermic/hypometabolic state (QIH) (*30*). This approach enables the direct investigation of how systemic metabolic demands and thermal states influence reproductive processes, including pregnancy (*31*); however, its direct impact on SEs^kiss^ remains elusive.

In the current study, we aimed to dissect the contributions of metabolic and thermal factors to reproductive suppression by examining their effects on SEs^kiss^ using chronic fiber photometry of ARC^kiss^ neurons.

## Results

### SEs^kiss^ are mildly suppressed by food restriction

To investigate the effect of negative energy balance on SEs^kiss^, we performed fiber photometry imaging of ARC^kiss^ neurons under food restriction (FR; 2.5 g of food provided at 24-h intervals). This represents 70–80% of the regular daily food intake (*23*) and has previously been used to prevent body weight gain and delay puberty onset in female mice (*5*). Both male and female mice were gonadectomized to remove gonadal negative feedback on ARC^kiss^ neurons, thereby facilitating SEs^kiss^ occurrence and amplitude (*18*, *20*, *21*). This enables the detection of potential effects of negative energy balance on SEs^kiss^. A Cre-dependent adeno-associated virus (AAV) driving GCaMP6s was injected into the ARC of *Kiss1-Cre* mice. We conducted photometry recordings at three time points, each for 24 h: (i) during FR, (ii) after the transition from FR to ad libitum feeding (FR-AL), and (iii) at least one day after the AL transition (post-AL) (Fig. 1A). Throughout the recording sessions, the core body temperature (Tc) was monitored using an implanted temperature logger. Post hoc histochemical analysis confirmed fiber placement and the efficiency and specificity of GCaMP6s expression (Fig. S1A and S1B).

**Fig. 1:**
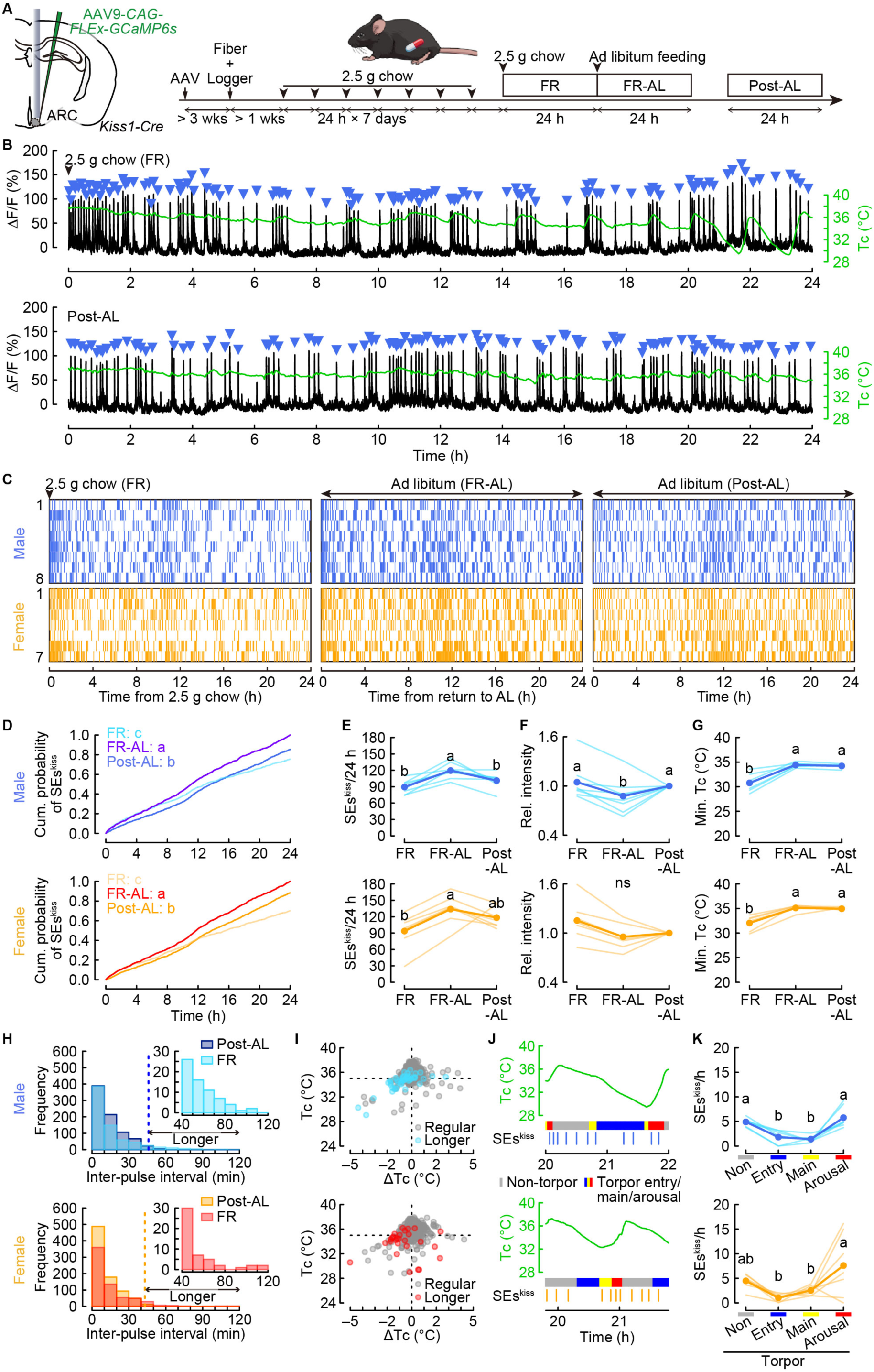
SEs^kiss^ are suppressed during daily torpor induced by food restriction. (A) Schematic of the experimental design and timeline. (B) Representative 24-h photometry traces (black) showing SEs^kiss^ (blue arrowheads) and Tc (green) under FR (top) and post-AL (bottom) conditions in castrated male mice. (C) Raster plots of SEs^kiss^ (vertical bars) in individual mice under FR (left), FR-AL (middle), and post-AL (right) conditions. (D–F) Cumulative probability (D), number (E), and relative (rel) intensity, expressed as normalized ΔF/F (F), of SEs^kiss^ in male (top) and female (bottom) mice. Different letters (a–c) indicate significant differences at *p* < 0.05 using the Kolmogorov–Smirnov test with Bonferroni correction (D). *n* = 8 castrated males; *n* = 7 ovariectomized females (C–F). (G) Minimum Tc during photometry recordings (*n* = 7 each for castrated males and ovariectomized females). Different letters (a, b) indicate significant differences at *p* < 0.05 using one-way repeated-measures ANOVA followed by Bonferroni-corrected post hoc tests (E–G). ns, not significant. (H) Histogram of inter-pulse intervals under FR and post-AL conditions. Dotted lines at 46 min (males) and 43 min (females) indicate the 99th percentile of inter-pulse intervals. Under FR, intervals are categorized as regular or prolonged based on this threshold (I) Scatter plots of ΔTc from the preceding SEs^kiss^ versus Tc at the current SEs^kiss^ under FR. Black and blue/red points denote regular and prolonged inter-pulse intervals, respectively. (J) Representative alignment of Tc (green), torpor phases (non-torpor, gray; entry, blue; maintenance, yellow; arousal, red), and SEs^kiss^ raster under FR. (K) Number of SEs^kiss^ across torpor phases under FR. Different letters (a, b) indicate significant differences at *p* < 0.05 using one-way repeated-measures ANOVA with Bonferroni-corrected post hoc tests. For more data, see Figs. S1 and S2.

Under chronic FR conditions, we observed sharp pulsatile ARC^kiss^ activity, closely resembling SEs^kiss^ previously reported in castrated (*20*) and ovariectomized (*21*) mice (Fig. 1B). SEs^kiss^ occurred most frequently soon after the supply of 2.5 g chow (time 0) and became sparser at later time points, as represented by raster plots across animals (Fig. 1C). After switching from FR to AL, the SEs^kiss^ frequency increased slightly but significantly in both sexes (Fig. 1D and 1E). Concomitant with this increase, SEs^kiss^ intensity was reduced by 12% in males, whereas the change in females was not statistically significant (Fig. 1F). We also noted the frequent clustering of SEs^kiss^ around 12 h after the onset of photometry, corresponding to the light-dark transition. This pattern persisted under post-AL conditions (Fig. 1C), with a significantly higher frequency immediately after the light-dark transition in both sexes (Fig. S1C and S1D). Overall, the SEs^kiss^ frequency and intensity were comparable between the FR and post-AL conditions (Fig. 1D–1F). Compared to the pronounced negative effects of FR on SEs^kiss^ and the marked rebound increase in activity following the transition to AL in peripubertal female mice (*5*), these data indicate that a chronic energy shortage has only minimal effects on SEs^kiss^ in gonadectomized adult mice.

Our previous study showed that under the same negative energy balance during the peripubertal period, hunger-responding ARC^Agrp^ neurons suppressed SEs^kiss^ frequency (*5*). However, in castrated adult male mice, chemogenetic activation of ARC^Agrp^ neurons after FR removal had no discernible effect on SEs^kiss^ frequency (Fig. S2). Together with the minimal impact of FR on SEs^kiss^ frequency in adults, these data suggest that SEs^kiss^ frequency is less influenced by ARC^Agrp^ neuron-mediated suppression and is robustly maintained despite a negative energy balance once sexual maturation occurs, probably because of metabolic differences in muscle and fat mass across life stages and gonadal states.

We noted that the FR condition induced daily torpor (Fig. 1B, 1G), consistent with previous studies (*29*, *32*). Despite the overall modest effects of FR on SEs^kiss^, raster plots and cumulative probability analysis revealed nocturnal suppression of SEs^kiss^ (Fig. 1C, 1D), suggesting inhibition by a caloric deficit or hypothermia during daily torpor. To examine these factors, we analyzed the relationship between SEs^kiss^ and Tc. Under post-AL conditions, most SEs^kiss^ inter-pulse intervals were clustered within 46 and 43 min in males and females, respectively. In contrast, longer intervals emerged under FR conditions for both sexes (Fig. 1H). These prolonged intervals occurred predominantly when the body temperature fell below 35 °C and declined steeply (Fig. 1I). Daily torpor can be divided into entry, maintenance, and arousal phases (*33*). Phase-based analysis showed that the SEs^kiss^ frequency was suppressed during torpor entry and maintenance (Fig. 1J, 1K), whereas SEs^kiss^ frequency during non-torpor and arousal periods (∼5 per hour) matched that observed under post-AL conditions. These data suggest that nocturnal suppression of SEs^kiss^ by food restriction is potentially linked to active hypothermia.

To further examine the relationship between daily torpor and SEs^kiss^ suppression, we performed fiber photometry over moderate (24 h; Fig. S3) and severe (48 h; Fig. 2A, 2B) fasting conditions. Three 24-h time windows were analyzed: (i) the first or second 24 h of fasting (Fasted), (ii) after returning from fasting to ad libitum feeding (Re-fed), and (iii) at least one day after re-feeding (Fed-post) (Fig. 2A). The temporal dynamics of SEs^kiss^ during the Fasted condition differed markedly from those under the Re-fed and Fed-post conditions (Fig. 2C, 2D; Figs. S3C, S3D, S4). The total number of SEs^kiss^ per 24 h was significantly reduced in the Fasted condition compared to the Fed-post condition (Fig. 2E; Fig. S3E), unlike the FR condition (Fig. 1E; Fig. S4J). Under the Re-fed condition, the SEs^kiss^ frequency recovered significantly (Fig. 2E; Fig. S3E), accompanied by a 15% reduction in SEs^kiss^ intensity in males, whereas the change in females was less pronounced (Fig. 2F; Fig. S3F). Consistent with a previous report (*28*), we observed clear daily torpor during fasting (Fig. 2G; Fig. S3G). Longer inter-pulse intervals of SEs^kiss^ emerged and occurred predominantly when body temperature fell below 35 °C and declined steeply (Fig. 2H, 2I; Fig. S3H, S3I). As in FR, phase-based torpor analysis showed that the SEs^kiss^ frequency was suppressed during torpor entry and maintenance (Fig. 2J, 2K; Fig. S3J, S3K). Notably, the SEs^kiss^ frequency during the torpor arousal phase (∼4 per hour) was comparable to that under post-AL conditions, despite a severe negative energy balance, supporting the notion that active hypometabolism or hypothermia, rather than energy scarcity per se, underlies SEs^kiss^ suppression.

**Fig. 2:**
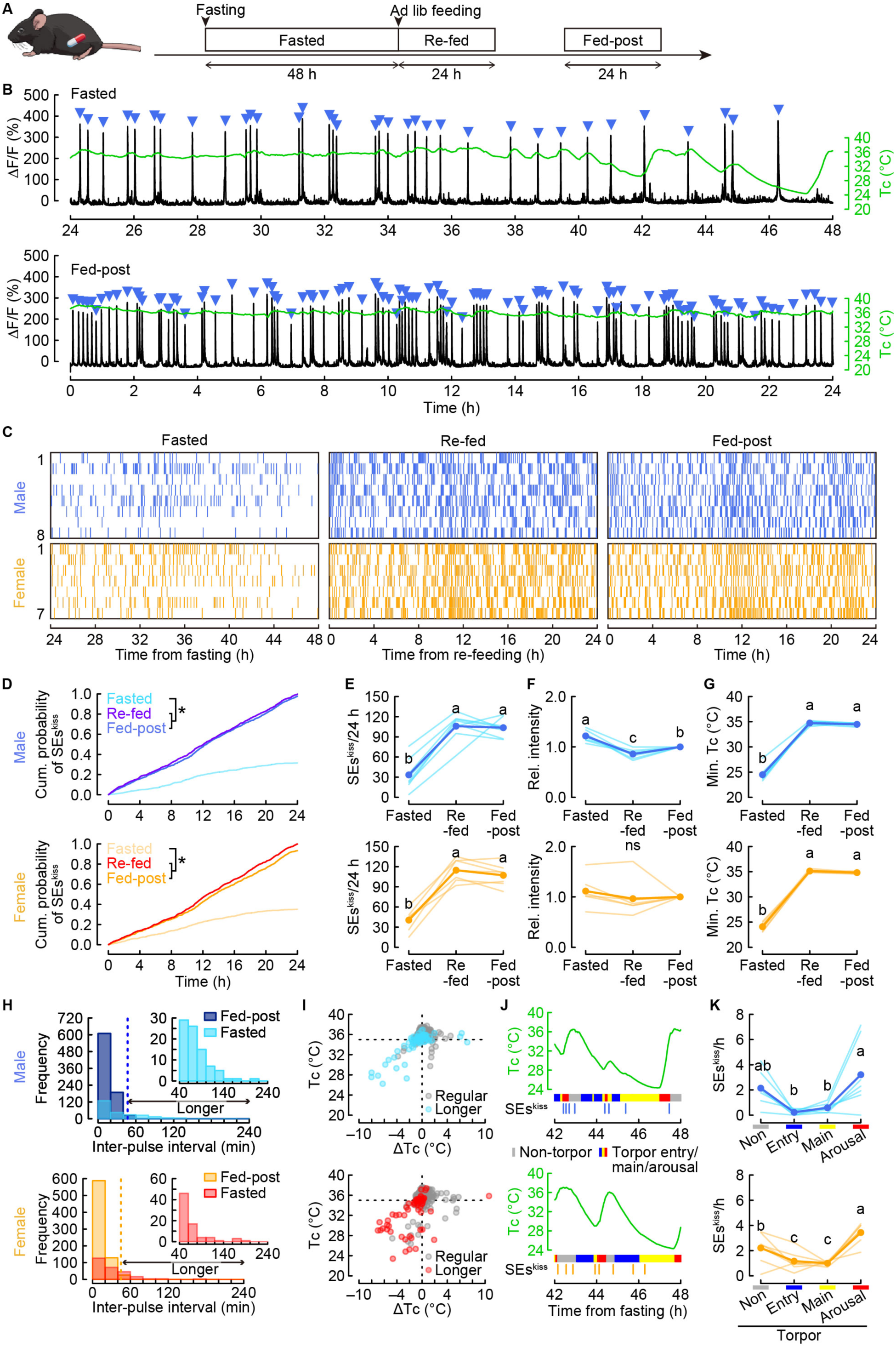
SEs^kiss^ are suppressed during deep torpor induced by 48-h fasting. (A) Schematic of the experimental timeline. (B) Representative 24-h photometry traces (black) showing SEs^kiss^ (blue arrowheads) and Tc (green) under 48-h Fasted (24–48 h window; top) and Fed-post (bottom) conditions in castrated male mice. (C) Raster plots of SEs^kiss^ (vertical bars) under Fasted (left; 24–48 h window), Re-fed (middle), and Fed-post (right) conditions. (D–F) Cumulative probability (D), number (E), and relative (rel) intensity, expressed as normalized ΔF/F (F), of SEs^kiss^. \**p* < 0.05 using the Kolmogorov–Smirnov test with Bonferroni correction (D). *n* = 8 castrated males; *n* = 7 ovariectomized females (C–F). (G) Minimum Tc during photometry recordings (*n* = 7 each). Different letters (a, b) indicate significant differences at *p* < 0.05 using one-way repeated-measures ANOVA with Bonferroni-corrected post hoc tests (E–G). ns, not significant. (H) Histogram of inter-pulse intervals under 48-h Fasted and Fed-post conditions. Dotted lines at 47 min (males) and 45 min (females) indicate the 99th percentile of inter-pulse intervals. Under the Fasted condition, intervals are categorized as regular or prolonged based on this threshold. (I) Scatter plots of ΔTc from the preceding SEs^kiss^ versus Tc at the current SEs^kiss^ in the Fasted group. Black and blue/red points denote regular and prolonged inter-pulse intervals, respectively. (J) Representative alignment of Tc (green), torpor phases (non-torpor, gray; entry, blue; maintenance, yellow; arousal, red), and SEs^kiss^ in the Fasted group. (K) Number of SEs^kiss^ across torpor phases in the Fasted group. Different letters (a–c) indicate significant differences at *p* < 0.05 using one-way repeated-measures ANOVA with Bonferroni-corrected post hoc tests. For more data, see Figs. S3 and S4.

### Pharmacological hypothermia suppresses SEs^kiss^

To further disentangle the metabolic and thermal contributions to SEs^kiss^ suppression, we examined the pharmacological effects of hypometabolism and hypothermia on SEs^kiss^. Systemic hypometabolism can be mimicked by the administration of 2-deoxy-D-glucose (2-DG), a glucose analog that inhibits glycolysis and disrupts glucose utilization, resulting in a dose-dependent decrease in the metabolic rate (*34*). A low dose of 2-DG (400 mg/kg) minimally affected Tc and blood glucose levels (BGLs), whereas a higher dose (1500 mg/kg) produced a modest decrease in Tc and larger changes in BGLs in both sexes (Fig. S5A–S5E). In contrast, N6-cyclohexyladenosine (CHA), a potent A1 adenosine receptor agonist known to induce hypothermia (*33*, *35*), markedly lowered Tc with minimal effects on BGLs (Fig. S5A–S5E).

Low-dose 2-DG and CHA produced comparable BGLs, but markedly different Tc values (Fig. S5A–S5E); therefore, we compared SEs^kiss^ dynamics after the administration of these agents. SEs^kiss^ were not suppressed by low-dose 2-DG, whereas CHA markedly suppressed SEs^kiss^ when Tc decreased substantially (Fig. 3A–3E). Phase-based analysis, analogous to torpor analysis, showed that SEs^kiss^ was completely abolished during entry into and maintenance of low Tc (Fig. 3F, 3G) and recovered ∼3.5 h after administration during the arousal phase (Fig. 3H) in both sexes. Females recovered SEs^kiss^ at a lower Tc than males (Fig. 3H), suggesting sex differences in the thermal threshold that permits SEs^kiss^. Notably, high-dose 2-DG caused significant changes in BGL (Fig. S5A–S5E) produced only a modest reduction in SEs^kiss^, which was much weaker than the effect of CHA (Fig. S5F–S5H). Collectively, these data indicate that active hypothermia, rather than hypometabolism, drives SEs^kiss^ suppression.

**Fig. 3:**
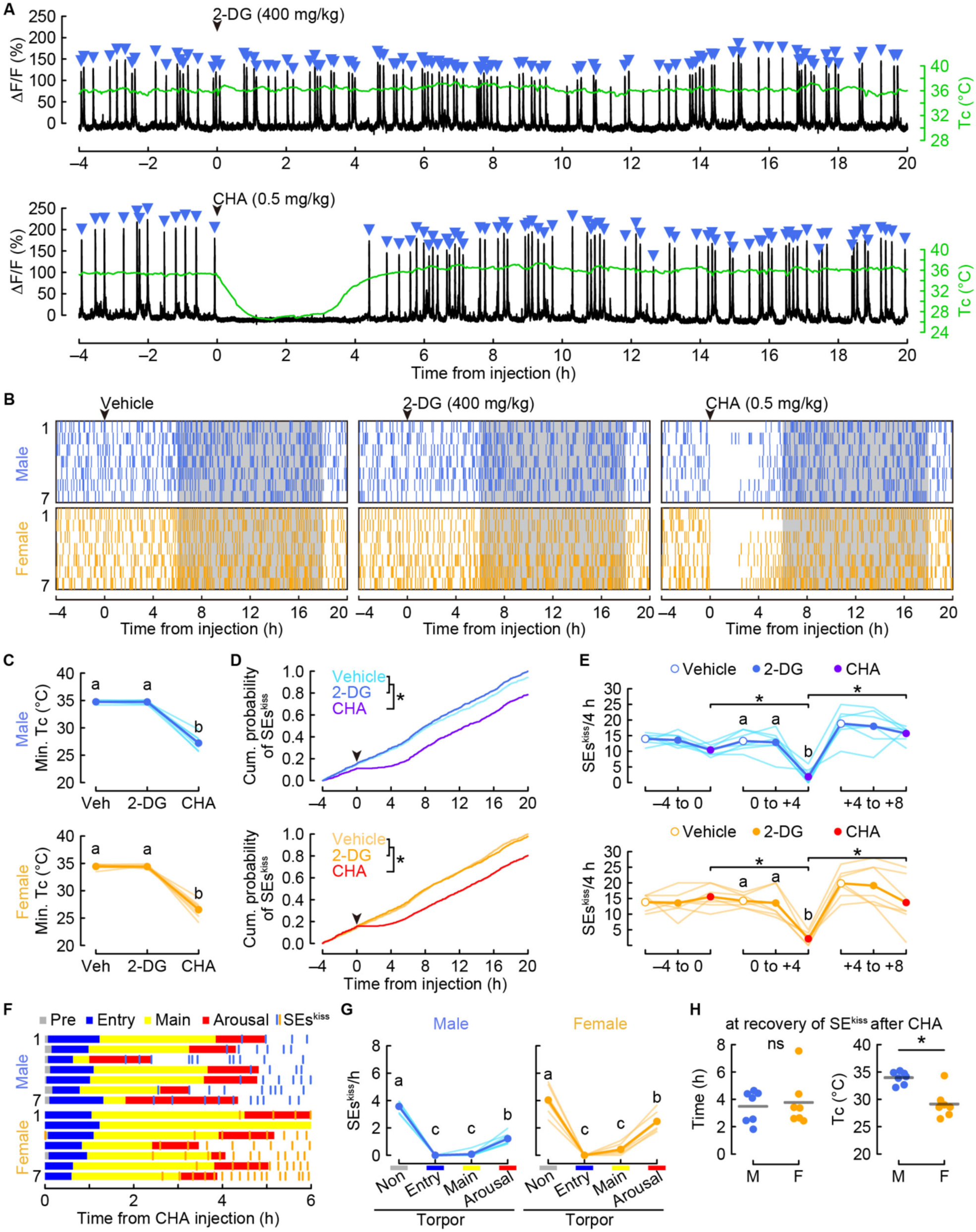
Pharmacological hypothermia suppresses SEs^kiss^. (A) Representative 24-h photometry traces (black) showing SEs^kiss^ (blue arrowheads) and Tc (green) in castrated male mice after 2-DG (400 mg/kg; top) or CHA (0.5 mg/kg; bottom) injection. Black arrowheads indicate the timing of intraperitoneal administration. (B) Raster plots of SEs^kiss^ (vertical bars) after vehicle (left), 2-DG (middle), and CHA (right) injection. (C) Minimum Tc during photometry recordings. (D) Cumulative probability of SEs^kiss^ over 24 h after vehicle, 2-DG, or CHA injection. \**p* < 0.05 using the Kolmogorov–Smirnov test with Bonferroni correction. (E) Number of SEs^kiss^ per 4-h time window after vehicle, 2-DG, or CHA injection. Different letters (a, b) indicate significant differences at *p* < 0.05 using one-way repeated-measures ANOVA with Bonferroni-corrected post hoc tests (C, E). (F) Torpor-phase plot (pre-torpor, gray; entry, blue; maintenance, yellow; arousal, red) aligned with SEs^kiss^ raster following CHA administration. (G) Number of SEs^kiss^ across torpor phases in the CHA group. Different letters (a–c) indicate significant differences at *p* < 0.05 using one-way repeated-measures ANOVA with Bonferroni-corrected post hoc tests. (H) Time to recovery (left) and Tc at recovery (right) of SEs^kiss^ after CHA administration. M, male; F, female. \**p* < 0.05 using two-sided non-paired *t*-test; ns, not significant. *n* = 7 each for castrated males and ovariectomized females (B–H). For more data, see Fig. S5.

### oQIH-mediated hypothermia suppresses SEs^kiss^

Hypothalamic Qrfp neurons can induce a hibernation-like hypothermic/hypometabolic state, termed QIH (*30*, *36*), providing a useful model for dissecting the active hypothermia of the reproductive axis. To obtain genetic access to ARC^kiss^ neurons independent of *Qrfp-iCre*, we generated *Kiss1-Flpo* knock-in mice using CRISPR-mediated genome editing (*37*) (Fig. S6A– S6D). To validate this line for fiber photometry-based Ca^2+^ imaging of ARC^kiss^ neurons, a Flpo-dependent AAV driving GCaMP6s was injected into the ARC of gonadectomized *Kiss1-Flpo* male and female mice (Fig. S6E). Post-hoc histochemical analysis showed that the targeting specificity and efficiency of *Kiss1-Flpo* were comparable to those of *the Kiss1-Cre* line (Figs. S1B, S6F, S6G). This efficiency was sufficient to reliably detect SEs^kiss^ events (Fig. S6H). The number of SEs^kiss^ was comparable between Cre- and Flpo-based fiber photometric recordings in both sexes during 24-h recordings under ad libitum feeding and during 48-h fasting (Fig. S6I). These data establish that the *Kiss1-Flpo* line is suitable for Ca^2+^ imaging in SEs^kiss^.

To examine the impact of optogenetically induced QIH (oQIH) on SEs^kiss^, we performed Ca^2+^ imaging in *Qrfp-iCre*; *Kiss1-Flpo* mice (Fig. 4A). hChR2(H134R) was selectively expressed in the anteroventral periventricular nucleus (AVPe) Qrfp neurons, and GCaMP6s was selectively expressed in ARC^kiss^ neurons (Fig. 4B). We confirmed robust oQIH induction in our preparation (movie S1), producing hypothermia with Tc approaching ambient temperature (∼25 °C), consistent with previous reports (*36*). Representative photometry traces, raster plots, and cumulative probability analysis revealed a marked suppression of SEs^kiss^ frequency during oQIH (Fig. 4C–4E). These data suggest that SEs^kiss^ are inhibited by the hypothermic state itself, hypometabolism, and/or by AVPe Qrfp neuronal input to ARC^kiss^ neurons. To distinguish between hypothermic and hypometabolic factors, we monitored SEs^kiss^ during oQIH while maintaining normothermia using a temperature-control device (warm-oQIH) (*38*). Under warm-oQIH, Tc was stably maintained near 37 °C in both sexes (Fig. S7A, top, movie S2). During hypothermic oQIH, the BGL transiently increased at the onset and offset, whereas during warm-oQIH, the BGL increase was attenuated, and the offset-associated increase occurred earlier (Fig. S7A, bottom). Notably, SEs^kiss^ frequency was fully restored during warm-oQIH to levels comparable to those observed in the pre- and post-oQIH periods (Fig. 4C–4E). This effect was not attributable to warmth-responsive neural mechanisms, as control mice expressing EYFP instead of ChR2 in Qrfp neurons (no QIH) on the temperature-control device showed no change in SEs^kiss^ frequency (Fig. S7B–S7D). Collectively, these data demonstrate that active hypothermia during QIH, rather than hypometabolism accompanying QIH, suppresses SEs^kiss^.

**Fig. 4:**
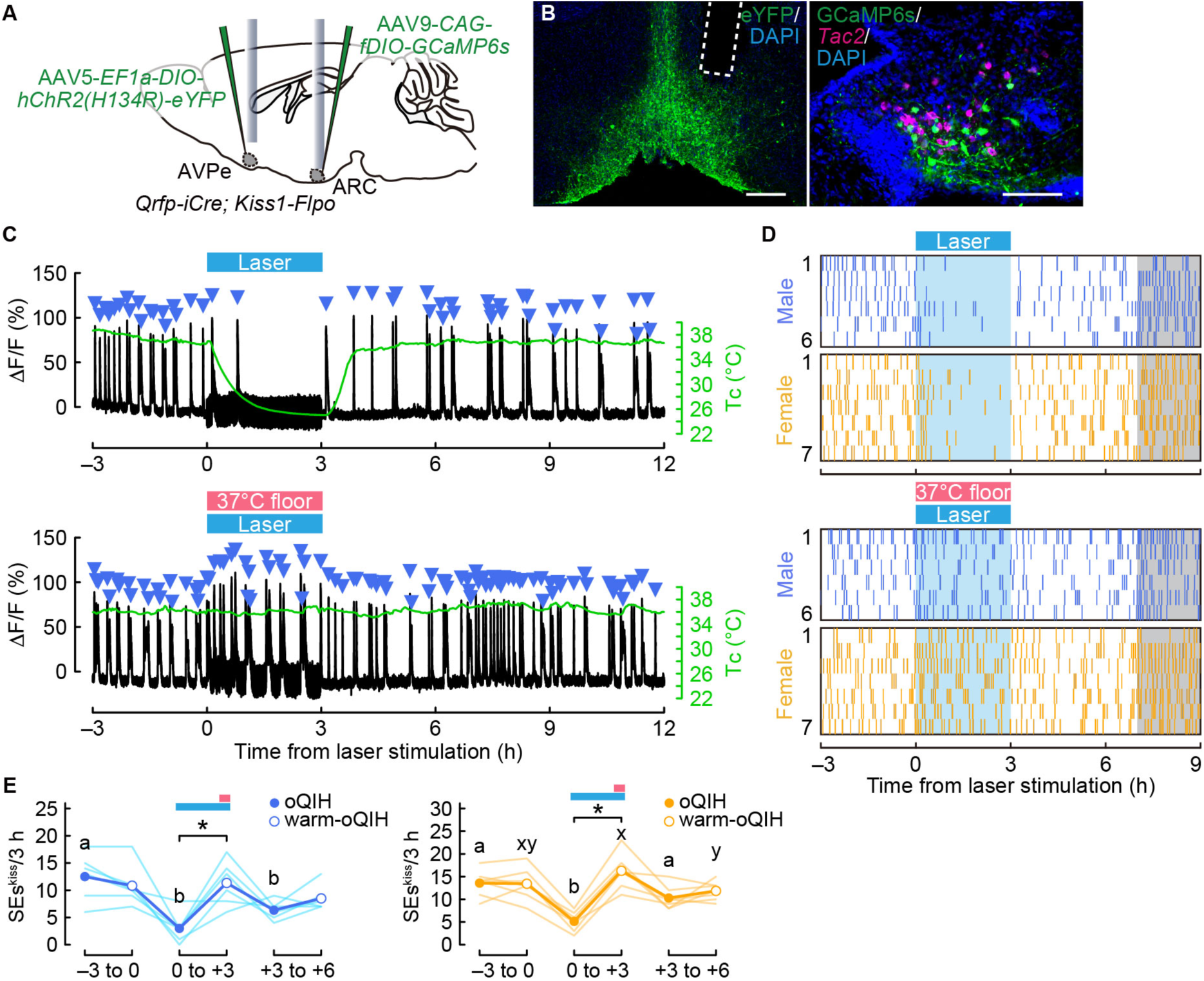
SEs^kiss^ are inhibited by oQIH-mediated hypothermia. (A) Schematic of the experimental setup. (B) Representative coronal sections of the AVPe (left), showing the optic fiber tract with hChR2(H134R)-EYFP expression, and the ARC (right), showing *Tac2* mRNA (magenta; a marker of ARC^kiss^ neurons) and GCaMP6s (green; anti-GFP) counterstained with DAPI (blue) from *Qrfp-iCre*; *Kiss1-Flpo* mice. Dashed line indicates fiber tract. Scale bars, 250 µm (left), 100 μm (right). (C) Representative 24-h photometry traces (black) showing SEs^kiss^ (blue arrowheads) and Tc (green) under oQIH (top) and warm-oQIH (bottom) conditions in castrated male mice. Blue and pink bars indicate the timing of optogenetic stimulation and floor heating (37 °C), respectively. (D) Raster plots of SEs^kiss^ (vertical bars) under oQIH (top) and warm-oQIH (bottom). (E) Number of SEs^kiss^ per 3-h time window in males (left) and females (right) under oQIH and warm-oQIH. Two-way repeated-measures ANOVA: QIH condition effect, *p* < 0.10; time effect, *p* < 0.05; interaction, *p* < 0.01 in males; QIH condition effect, *p* < 0.01; time effect, *p* < 0.05; interaction, *p* < 0.01 in females. \**p* < 0.01 by post hoc two-tailed paired *t*-test. Different letters (a, b for oQIH; x, y for warm-oQIH) indicate significant differences at *p* < 0.05 using one-way repeated-measures ANOVA with Bonferroni-corrected post hoc tests within each condition. *n* = 6 castrated males; *n* = 7 ovariectomized females (D, E). For more data, see Figs. S6 and S7.

### Long-term hypothermia inhibits gametogenesis

Lastly, we examined the chronic impact of hypothermia on reproductive function. We confirmed that chronic SEs^kiss^ suppression corresponded to a hypothermic state during chemogenetically induced QIH (Fig. S8). We assessed gonadal phenotypes after repeated chemogenetically induced QIH every 4–5 days for four weeks (Fig. 5A; see Methods), a duration sufficient to disrupt cyclic follicular development occurring in the 4–6-day estrus cycle and to affect spermatogenesis, given the ∼35-day seminiferous epithelial cycle in mice. Post hoc histology confirmed the targeted hM3Dq-mCherry expression in the AVPe (Fig. 5B). Implanted temperature loggers verified robust clozapine-N-oxide (CNO)-induced QIH in the hM3Dq group. In contrast, the mCherry controls maintained a stable Tc near 37 °C (Fig. 5C). Body weight remained unchanged after repeated QIH administration (Fig. 5D). In males, however, the hM3Dq group exhibited significantly reduced testicular weight compared to the mCherry controls (Fig. 5E), along with a marked depletion of spermatocytes and spermatids (Fig. 5F, 5G), indicating that hypothermia impairs spermatogenesis. Similarly, in females, the hM3Dq group showed significantly reduced ovarian weights compared with the mCherry controls (Fig. 5H), along with a marked decrease in the number of antral follicles and corpora lutea (Fig. 5I, 5 J), indicating that hypothermia arrests the follicular development. Collectively, these data demonstrate that repeated hypothermia associated with SEs^kiss^ suppression leads to a pronounced decline in the reproductive capacity of both sexes.

**Fig. 5:**
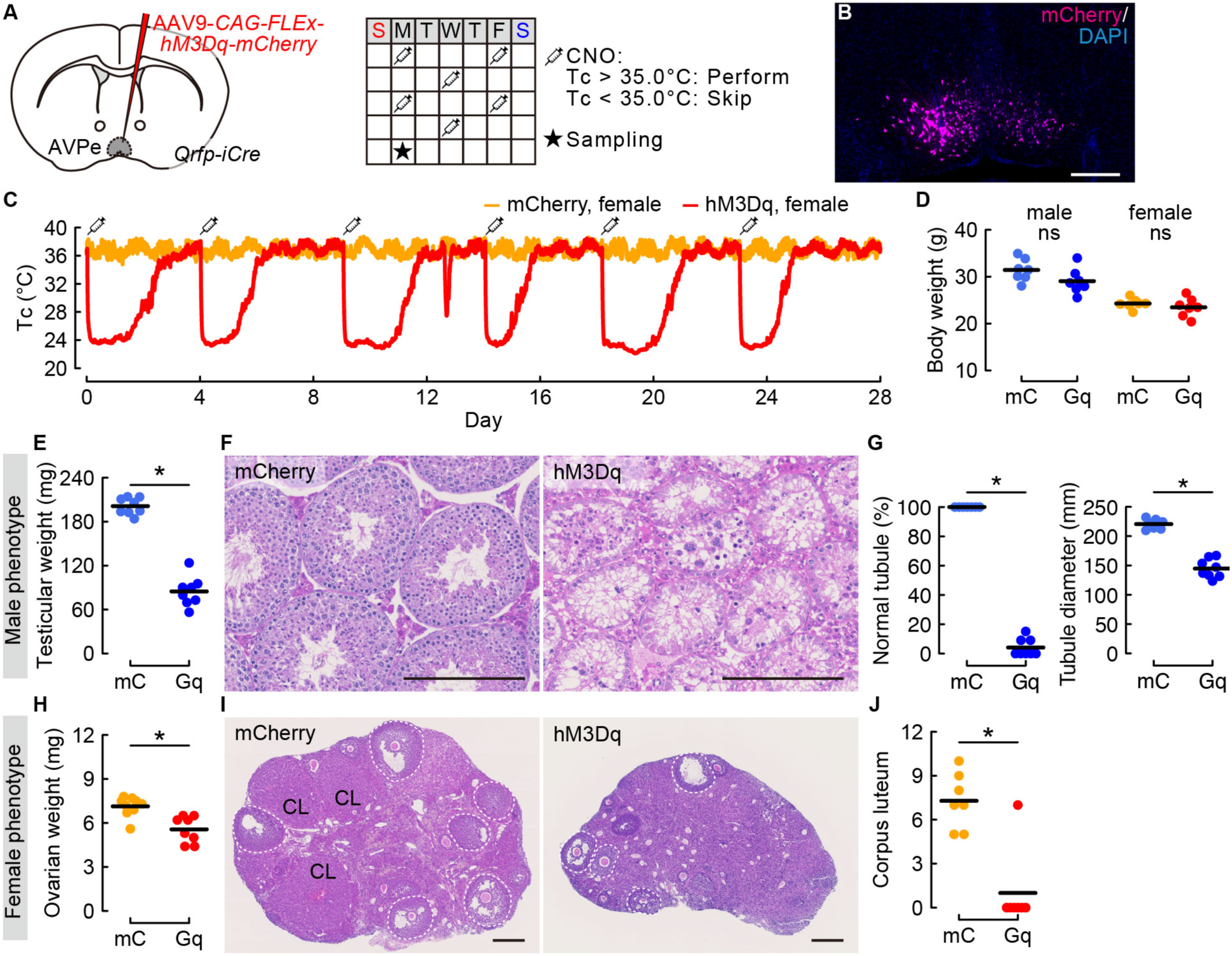
Repeated QIH impairs gametogenesis. (A) Schematic of the experimental setup (left) and an example of CNO injection timeline. (B) Representative coronal sections of the AVPe with hM3Dq-mCherry expression, counterstained with DAPI (blue). Scale bar, 250 μm. (C) Representative Tc traces over 4 weeks. Syringes indicate the timing of CNO injection. (D) Body weight at the end of the CNO injection period in mCherry control (green) and hM3Dq (blue) males, and mCherry control (orange) and hM3Dq (red) females. (E) Testicular weights after CNO treatment. (F) Representative hematoxylin and eosin-stained testicular sections. Scale bars, 200 μm. (G) Rate of normal seminiferous tubule (left) and tubule diameter (right) after CNO treatment. (H) Ovarian weights after CNO treatment. (I) Representative hematoxylin and eosin-stained ovarian sections. CL, corpus luteum. Scale bars, 200 μm. (J) Number of corpora lutea after CNO treatment. \**p* < 0.05 using the Mann–Whitney U test. *n* = 7 per group (E, G, H, J). For more data, see Fig. S8.

## Discussion

Organisms have evolved stringent checkpoints to regulate energy allocation to reproductive functions as an adaptive strategy for survival in harsh environments. In the wild, many mammals strategically suspend reproduction during winter or periods of resource scarcity. However, our understanding of the contributions of metabolic and thermal factors to reproductive suppression is limited. In this study, we demonstrate that active hypothermia under low-energy conditions is the primary driver of reduced SEs^kiss^ frequency, leading to impaired GnRH pulse generation and suppression of gametogenesis. This study provides insight into how thermal signals act as critical gatekeepers of the mammalian reproductive system. Here, we discuss the biological insights obtained from the current study and its limitations.

We show that the SEs^kiss^ frequency is suppressed by diet-, pharmacologically, and neuronally induced hypothermia. An important question is whether hypothermia selectively suppresses ARC^kiss^ neural activity or broadly reduces neural activity throughout the nervous system. While SEs^kiss^ are markedly suppressed during QIH at a Tc of approximately 25 °C, previous ex vivo studies have shown that circadian rhythms of clock gene transcription and intracellular Ca^2+^ transients in the suprachiasmatic nucleus (SCN), the master circadian clock, are maintained even at 22 °C (*39*). Whereas neural activity in the hippocampus CA1 is profoundly suppressed during QIH (*40*), vesicular glutamate transporter type 3-expressing neurons in the medullary raphe pallidus are active immediately after the arousal phase during the QIH state, well before the Tc returns to the normothermic range (*41*). At the circuit level, sensory responsiveness is substantially reduced during deep torpor but is not completely blocked; sufficiently strong stimuli can trigger arousal (*42*). These observations suggest the presence of cell type-specific mechanisms of neural suppression during hypothermia. In addition to circuit-level inhibition, temperature-dependent biochemical reactions of intracellular proteins may have contributed to the suppression of SEs^kiss^.

We demonstrate that fasting impairs SEs^kiss^ frequency, extending previous findings that fasting suppresses *Kiss1* gene expression (*43*, *44*) and pulsatile luteinizing hormone (LH) secretion (*7*, *45–49*). In the context of reproductive suppression under malnutrition (*50–52*), acute glucose deprivation induced by the central (*53*, *54*) and peripheral (*54–56*) administration of 2-DG has been shown to suppress pulsatile LH secretion downstream of SEs^kiss^ (*18*, *20*, *21*). The present study updates this view by suggesting that 2-DG-induced suppression of LH pulses is primarily caused by subsequent hypothermia rather than glucose deprivation. Systemic glucose deprivation activates PVH (*9*) and SON neurons that express dynorphin (*54*), potentially inhibiting ARC^kiss^ neural activity. Elucidating the mechanisms underlying these neural circuits under conditions of glucose deprivation and hypothermia is an important direction for future research. The hypothermic effects of the potent adenosine A1 receptor agonist, CHA, involve both the peripheral and central pathways (*57*), which are thought to be mediated by A1 receptors in the dorsomedial hypothalamus (*57*) or the nucleus of the solitary tract (*58*). Direct manipulation of the downstream targets of A1 receptor-expressing neurons would help clarify whether CHA-c induced SEs^kiss^ suppression is driven by hypothermia itself or specific neural inputs. Notably, our phase-based analyses indicate that the suppression of SEs^kiss^ frequency under food restriction, fasting, pharmacological intervention, and QIH is specific to the entry and maintenance phases but not to the arousal phase, even when Tc remains well below the normothermic range. These findings support the notion that hypothermia suppresses SEs^kiss^ via thermoregulatory neural inputs to the ARC^kiss^ neurons.

Our data indicate no apparent sex differences in SEs^kiss^ suppression under a negative energy balance, despite expectations based on known sex differences in basal metabolism (*59*). BGLs showed moderate sex differences upon pharmacological intervention and QIH. During QIH, we replicated the previously reported BGL dynamics, with an acute increase followed by a decline (*60*). The physiological significance of the rebound increase in BGLs after QIH remains to be determined.

We acknowledge several limitations of the present study. First, we performed SEs^kiss^ monitoring in gonadectomized mice to facilitate its detection. Because sex steroid-mediated negative feedback contributes to the suppression of LH pulses induced by fasting (*61*) or 2-DG (*62*) in rats, the discrepancies between previous studies (*5*, *24*) and the current study may be attributable to the use of gonadectomized animals. In this context, SEs^kiss^ is released from gonadal negative feedback and remains elevated, potentially masking the relatively subtle inhibitory regulatory effects. Second, we did not examine the reversibility of gametogenesis following suppression by repeated QIH. It would be important in future studies to determine how repeated QIH affects estrus cyclicity and the maturation of early-stage germ cells, including primordial and primary follicles and spermatogonia. Third, although we measured the BGLs during hypothermia, we did not perform a comprehensive metabolic assessment. Thyroid hormones, implicated in seasonal reproduction (*63*) and hibernation (*64*), are potential candidates for future investigations. More broadly, because only a limited number of circulating metabolic factors can currently be measured with high temporal resolution, future studies should expand the use of photometry-compatible biosensors that convert ligand binding into fluorescent signals, such as GPCR activation-based sensors (*65*). In parallel, advances in wearable measurement technologies, potentially enabled by engineering approaches, such as needle-shaped diamond electrodes (*66*), may allow more comprehensive, real-time monitoring of physiological states.

## Materials and Methods

All animal procedures were approved by the Institutional Animal Care and Use Committee of the RIKEN Kobe Branch. C57BL/6N female mice were purchased from Japan SLC, Inc. (Shizuoka, Japan). *Kiss1-Cre* (JAX #017701) mice were obtained from The Jackson Laboratory. *Qrfp-iCre* mice have been previously described (*30*). *Kiss1-Flpo* mice were generated in this study (see below for further details). Animals were housed at the RIKEN Center for Biosystems Dynamics Research animal facility under controlled ambient temperature (18–23 °C) and a 12-h light/12-h dark cycle. Unless otherwise specified, the mice had ad libitum access to standard laboratory chow (MFG; Oriental Yeast, Shiga, Japan; 3.57 kcal/g) and water.

### Generation of *Kiss1-Flpo* knock-in mice

The *Kiss1*-Flpo knock-in mouse line (Accession No. CDB0355E; https://large.riken.jp/distribution/mutant-list.html) was generated using CRISPR/Cas9-mediated knock-in in zygotes as previously described (*37*). The SV40 nuclear localization signal (NLS) was added to the 5′ of Flpo open reading frame by PCR primers 5′- GGCGCGCCACCATGGCTCCTAAGAAGAAGAGGAAGGTGATGAGCCAGTTCGACATC CTG; 5′-GTCGACTCAGATCCGCCTGTTGATGTAG. The plasmid *pTCAV-FLEx(loxP)-FlpO* (#67829; Addgene) was used as the PCR template. Then, a targeting vector comprising *Flpo-polyA* was constructed using the PCR-based method and inserted just after the start codon of *Kiss1* gene. Guide RNA (gRNA) sites were designed using CRISPRdirect (*67*) to target locations upstream and downstream of the start codon (Fig. S6). For microinjection, a mixture of three CRISPR RNAs (crRNAs) (50 ng/ µL), trans-activating crRNA (tracrRNA) (300 ng/µL), donor vector (10 ng/µL), and Cas9 protein (100 ng/µL) was injected into the pronucleus of a C57BL/6 one-cell stage zygote. Kiss1 crRNA#1 (5′-AGA AGC CAU UGA GAU CAU UCg uuu uag agc uau gcu guu uug), Kiss1 crRNA#2 (5′- CAG GUA CGC ACU GUC CGA UAg uuu uag agc uau gcu guu uug), PITCh 3 crRNA (5′-GCA UCG UAC GCG UAC GUG UUg uuu uag agc uau gcu guu uug), and tracrRNA (5′-AAA CAG CAU AGC AAG UUA AAA UAA GGC UAG UCC GUU AUC AAC UUG AAA AAG UGG CAC CGA GUC GGU GCU) were purchased from FASMAC (Atsugi, Japan). As a result, 38 F0 founder mice were obtained, five (four males and one female) of which were targeted as identified using 5′ and 3′ junction PCR. We further analyzed the 5′ and 3′ junctions and full-length sequencing from four males. The following primers were used for PCR and sequencing analyses:

For the detection of the Flpo internal sequence: *Flpo-F* 5′- CTGGCCACATTCATCAACTGCGG; *Flpo-R* 5′- CTTCTTCAGGGCCTTGTTGTAGCTG.

For the 5′ boundary: *Kiss1-F* 5′-CCTGGCTTCTGTACATCCTTCTGGCCATC; *Flpo-*5′*R* 5′-GTCTTGCACAGGATGTCGAACTGGCTC.

For the 3′ boundary: *FLPo-*3′*F* 5′-AGCATCAGATACCCCGCCTGGAACG; *Kiss1-R* 5′- CTAGCTGAGCCTCCAGTGCTCACAGC.

For the full-length: *Kiss1-F* 5′-CCTGGCTTCTGTACATCCTTCTGGCCATC; *Kiss1-R* 5′- CTAGCTGAGCCTCCAGTGCTCACAGC.

The PCR products using the primer sets were subcloned into the pCR Blunt II TOPO vector (Zero Blunt TOPO PCR Cloning Kit, Thermo Fisher Scientific) and sequenced using *M13-Forward* and *M13-Reverse* primers. Frameshift deletions were found in the allele opposite to the knock-in allele, resulting in *Kiss1* knockout and infertility. To overcome infertility in F0 founder males, spermatogenesis was supported by T supplementation for six weeks (*68*). F1 mice were produced by in vitro fertilization with sperm from T-supplemented mice and wild-type oocytes. *Kiss1-Flpo* allele was confirmed by genotyping F1 mice. The line was established using two males harboring a target sequence identified using PCR and sequencing. Genotyping PCR was performed using internal primers *Flpo-F* and *Flpo-R*, as described above (see Fig. S6 for details).

### Viral preparations

The following AAV vectors were purchased from Addgene. The titer is expressed as genomic particles (gp) per mL.

AAV serotype 9 *CAG-FLEx-GCaMP6s-WPRE-SV40* (1.7 × 10^13^ gp/mL, #100842-AAV9)

AAV serotype 5 *EF1a-double floxed-hChR2(H134R)-EYFP-WPRE-HGHpA* (1.8 × 10^13^ gp/mL, #20298-AAV5)

AAV serotype 5 *EF1a-DIO-EYFP-WPRE-HGHpA* (2.3 × 10^13^ gp/mL, #27056-AAV5)

AAV serotype 8 *hSyn-DIO-hM3D(Gq)-mCherry* (2.2 × 10^13^ gp/mL, #44361-AAV8)

AAV serotype 8 *hSyn-DIO-mCherry* (2.0 × 10^13^ gp/mL, #50459-AAV8)

The following AAV vectors were purchased from the Canadian Neurophotonics Platform Viral Vector Core Facility (RRID: SCR_016477).

AAV serotype 9 *CAG-fDIO-GCaMP6s* (7.5 × 10^12^ gp/mL, construct-1347-aav2-9)

AAV serotype 8 *hSyn-fDIO-hM3D(Gq)-mCherry* (2.0 × 10^13^ gp/mL, construct-1457-aav2-8)

### Stereotaxic injection

For AAV injection and optical fiber implantation, the mice were anesthetized with an intraperitoneal injection of saline containing 65 mg/kg ketamine (Daiichi Sankyo, Tokyo, Japan) and 13 mg/kg xylazine (X1251; Sigma-Aldrich), and head-fixed to a stereotaxic apparatus (#68045, RWD). Immediately after brain surgery, mice were gonadectomized. Photometric recordings were then performed after a recovery period of at least four weeks.

To perform the experiments in Figs. 1–3, S1, and S3–S5, the AAV carrying a Cre-dependent GCaMP6s (AAV9 *CAG-FLEx-GCaMP6s-WPRE-SV40*) was injected into the ARC of *Kiss1-Cre* female and male mice (4–7 months old) using the following coordinates from the bregma: posterior 1.95 mm, lateral 0.2 mm, and ventral 5.9 mm. In Fig. S2, a 1:3 mixture of the AAV carrying a Cre-dependent *GCaMP6s* and an AAV carrying a Flpo-dependent *hM3D(Gq)- mCherry* (AAV8 *hSyn-fDIO-hM3D(Gq)-mCherry*) was injected into the ARC of *Kiss1-Cre*; *Agrp-Flpo* male mice. In Fig. S6, the AAV carrying a Flpo-dependent *GCaMP6s* (AAV9 *CAG-fDIO-GCaMP6s*) was injected into the ARC of *Kiss1-Flpo* female and male mice (2–5 months-old). A total of 200 nL of AAV was injected into the ARC at a speed of 50 nL/min using a glass capillary regulated by a micro syringe pump injector (UMP3; World Precision Instruments, Sarasota, FL, USA). An optical fiber (NA = 0.50, core diameter = 400 µm from Kyocera) was placed above the ARC using the following coordinates from the bregma: posterior 1.95 mm, lateral 0.2 mm, and ventral 5.8 mm, defined on a brain atlas (*69*). The injected fibers were fixed to the skull using dental cement. Postsurgical mice were housed individually and allowed ad libitum access to food and water.

To perform the experiments in Fig. 4 and Fig. S7, the AAV carrying a Cre-dependent *hChR2-EYFP* or *EYFP* control and AAV carrying a Flpo-dependent *GCaMP6s* were injected into the AVPe and ARC of ovariectomized *Qrfp-iCre*; *Kiss1-Flpo* female and male mice (2–7 months old) using the following coordinates from the bregma: anterior 0.60 mm, bilateral 0.25 mm, and ventral 5.40 mm to the AVPe; and posterior 1.95 mm, lateral 0.2 mm, and ventral 5.9 mm to the ARC, respectively. A total of 300 and 200 nL of AAV were injected into the AVPe and ARC at a speed of 60 and 50 nL/min, respectively, using a glass capillary controlled by a micro syringe pump injector. An optical fiber was placed above the AVPe and ARC using the following coordinates from the bregma: anterior 0.60 mm, lateral 0.0 mm, and ventral 5.2 mm to the AVPe; and posterior 1.95 mm, lateral 0.2 mm, and ventral 5.8 mm to the ARC, respectively.

To perform the experiments in Fig. 5, the AAV carrying a Cre-dependent *hM3D(Gq)* was injected into the AVPe of gonadal-intact *Qrfp-iCre* female and male mice (3–5 months old) using the following coordinates from the bregma: anterior 0.60 mm, bilateral 0.25 mm, and ventral 5.40 mm to the AVPe. A total of 300 nL of AAV was injected into the AVPe at a speed of 60 nL/min using a glass capillary controlled by a micro syringe pump injector.

### Fiber photometry recording

Fluorescence signals were acquired using a fiber photometry system, based on a previously published design (*5*, *19*). All the optical components were purchased from Doric Lenses (Quebec, Canada). We performed chronic Ca^2+^ imaging by delivering an excitation light (470 nm modulated at 530.481 Hz) and collecting the emitted fluorescence using an integrated Fluorescence Mini Cube (Doric Lenses, iFMC4_IE(405)_E(460–490)_F(500–550)_S). Light collection, filtering, and demodulation were performed using the Doric Photometry Setup and Doric Neuroscience Studio Software (Doric Lenses). The 470-nm signal was recorded as calcium-dependent GCaMP6s. The power output at the fiber tip was approximately 0.5–5 µW. Signals were initially acquired at 12 kHz and then decimated to 120 Hz.

We used a custom R script for the analyses. Because the signal intensity of SEs^kiss^ varies widely due to variations in surgery, we first calculated the average ΔF/F height of the stereotyped SE^kiss^ peak for each mouse from several visually obvious peaks, using a full-width at half-maximum threshold over 10 seconds (*5*, *19*). The ΔF/F was calculated by (Ft – F_0_)/F_0_, where Ft is the recorded signal at time = t, and F_0_ is the average of signals over the entire 24 hours of recording. Then, SEs^kiss^ were automatically detected using the findpeaks function in R, with the peak height threshold as 0.4-fold of the average ΔF/F height.

### Core body temperature measurement

Core body temperature was recorded at 1-min interval using Anipill loggers (BodyCap, Paris, France) from freely moving mice. Logged pills were implanted into the intraperitoneal cavity under anesthesia. Gentamicin (5 mg/kg; Takata Pharmaceutical) was administered intraperitoneally immediately after the surgery to prevent infection. After 5–7 days of recovery following surgery, core body temperatures were recorded every 1 min using an AniLogger Monitor. Data were extracted using the CSV file and analyzed using R.

### Classification of torpor entry, maintenance, and arousal

We classified mice with core body temperatures of 35 °C or higher as a non-torpor state, and those with core body temperatures of 35 °C or lower as a torpor state, as defined in previous studies (*28*, *33*). If a mouse was in a torpor state and its core body temperature was decreasing at a rate of 0.05 °C/min or higher, that mouse was classified as a torpor entry. If a mouse was in a torpor state and its core body temperature was rising at 0.1 °C/min or faster, the mouse was classified as a torpor arousal. The other torpor states, excluding torpor entry and arousal, were classified as torpor maintenance states.

### Fasting and food restriction

The food was removed, and the bedding was renewed immediately before fiber photometric recording, as shown in Fig. 2 and Fig. S3. On the re-feeding day, sufficient food was supplied to the recording cage at the onset of the photometric recording.

Food was removed, and the bedding was renewed. The mice were then provided with 2.50 ± 0.01 grams of baby food (CLEA Japan Inc., Tokyo, Japan; 3.92 kcal/g) daily at Zeitgeber time 1–2 with ad libitum access to water. Mice received manually or automatically dispensed food at 24-h intervals via a customized feeding system based on an automatic pet feeder (Highshop, #FC0196200pufF) mounted on the cage lid (*5*).

### Optogenetic stimulation

The fiberoptic cannulas implanted in the mice were connected to a fiberoptic patch cable (200 µm diameter, NA: 0.22, 1.0 m length, Doric Lenses) using ceramic sleeves (Thorlabs) before fiber-photometric recording. Diode-pumped solid-state lasers (465 nm blue, IOS-465, RWD) were used for optogenetic stimulation. The laser output at the optical fiber tip was adjusted to 6–8 mW.

Laser stimulation was applied at 2 Hz with a 10-ms pulse width for 3 h to induce QIH (*41*). To induce warm-oQIH, the measurement cage was placed on a heating pad simultaneously with the laser stimulation.

### Chemogenetic manipulation

Clozapine N-oxide (CNO, #4936, Tocris Bioscience) was dissolved at 0.5 mg/mL for intraperitoneal injection. CNO was intraperitoneally injected at a dose of 2 mg/kg.

### Blood glucose monitoring

Blood glucose levels were monitored using a blood glucose meter for laboratory animals (SUGL-001; ForaCare Japan, Tokyo, Japan) after blood samples were collected from the tail. The animals were habituated for 2 days before measurements to prevent potential stress-induced elevation in blood glucose.

### Histochemistry

Mice were anesthetized with isoflurane and perfused with phosphate-buffered saline (PBS) followed by 4% paraformaldehyde (PFA) in PBS. The brains and ovaries were post-fixed overnight with 4% PFA in PBS. Coronal brain sections (30 µm; every fourth section) were prepared using a cryostat (Leica). To generate cRNA probes for *Tac2*, DNA templates were amplified from whole-brain cDNA by PCR (Genostaff, cat#MD-01). A T3 RNA polymerase recognition site (5′-AATTAACCCTCACTAAAGGG) was added to the 3′ end of the reverse primers. The forward (F) and reverse (R) primers to generate DNA templates for cRNA probes are as follows: F 5′- AGCCAGCTCCCTGATCCT; R 5′-TTGCTATGGGGTTGAGGC for *Tac2*. DNA templates for *Tac2* were subjected to in vitro transcription with DIG (cat#11277073910)- RNA labeling mix and T3 RNA polymerase (cat#11031163001), according to the manufacturer’s instructions (Roche Applied Science).

Fluorescent in situ hybridization (ISH) combined with anti-GFP immunohistochemical staining was performed as previously reported (ref). Briefly, after hybridization and washing, brain sections were incubated with a peroxidase (POD)-conjugated anti-fluorescein antibody (Roche Applied Science, cat#11426346910, 1:500) overnight at 4 °C. The following day, the signals were amplified with TSA-plus Cyanine 5 (NEL745001KT, Akoya Biosciences; 1:70 in 1×Plus amplification diluent) for 25 min. After washing with PBS containing 0.1% Tween-20 (PBST) for 5 min, POD was inactivated with 2% sodium azide in PBS for 15 min, followed by three washes with PBST for 10 min. Sections were then incubated overnight with horseradish peroxidase (HRP)-conjugated anti-DIG (Roche Applied Science cat #11207733910, 1:500) and anti-GFP (Aves Labs cat #GFP-1010, 1:200) antibodies. Signals were amplified using TSA-plus Cyanine 3 (NEL744001KT, Akoya Biosciences; 1:70 in 1×Plus amplification diluent) for 25 min, followed by washing, and GFP-positive cells were visualized using anti-chicken Alexa Fluor 488 (Jackson ImmunoResearch cat#703-545-155, 1:250). PBS containing 50 ng/mL 4’,6-diamidino-2-phenylindole dihydrochloride (DAPI; Sigma-Aldrich, cat#D8417) was used for nuclear staining. Images were acquired using an Olympus BX53 microscope equipped with a 10×(N.A. 0.4) objective lens. The cells were counted manually.

We compared *Tac2* gene expression in the ARC as a robust marker of ARC^kiss^ neurons because the *Kiss1* gene expression is not sufficient for stable detection using conventional ISH.

The ovaries were then embedded in paraffin. Serial 5-µm sections (every 10th) were prepared using a microtome. The paraffin sections were deparaffinized with xylene, hydrated with ethanol, and stained with hematoxylin and eosin for light microscopy. The corpus lutea were manually counted. Follicle size was quantified using Fiji software.

The testes were fixed in Bouin’s solution overnight and washed with 70% ethanol. These were then embedded in paraffin and sectioned to yield 3-μm thick sections. The paraffin sections were deparaffinized with xylene, hydrated with graded ethanol, and stained with hematoxylin and eosin for light microscopy. The stage of the seminal tubule was quantified using Fiji software.

### Statistical analysis

The statistical details of each experiment, including the statistical tests used, exact value of n, and what n represents, are described in each figure legend. R version 4.3.3 (http://www.R-project.org/) was used for all analyses. Statistical significance was set at *p* < 0.05.

## Acknowledgments

We thank the staff at the RIKEN Biosystems Dynamics Research (BDR) animal facility for animal care and in vitro fertilization; Yoshifumi Yamaguchi (Hokkaido University), Genshiro Sunagawa (RIKEN BDR), Hiroko Tsukamura (Nagoya University), and members of the Miyamichi laboratory for critically reading the manuscript; and the Canadian Neurophotonics Platform Viral Vector Core Facility for AAV production.

## Author contributions

Conceptualization: KM, TG

Methodology: MH, JH, TA, TS, TG

Investigation: MT, DS, TG

Supervision: KM, TG

Writing—original draft: MH, KM, TG

Writing—review & editing: KM, TG

## Data and materials availability

Original fiber photometry data and custom analysis scripts were deposited in the SSBD repository at https://doi.org/10.24631/ssbd.repos.2026.08.531. All other data are available in the main text and the Supplementary Materials. All materials, including *Kiss1-Flpo* mice, are available through requests to the corresponding author.

## Supplementary Figures

**Fig. S1:**
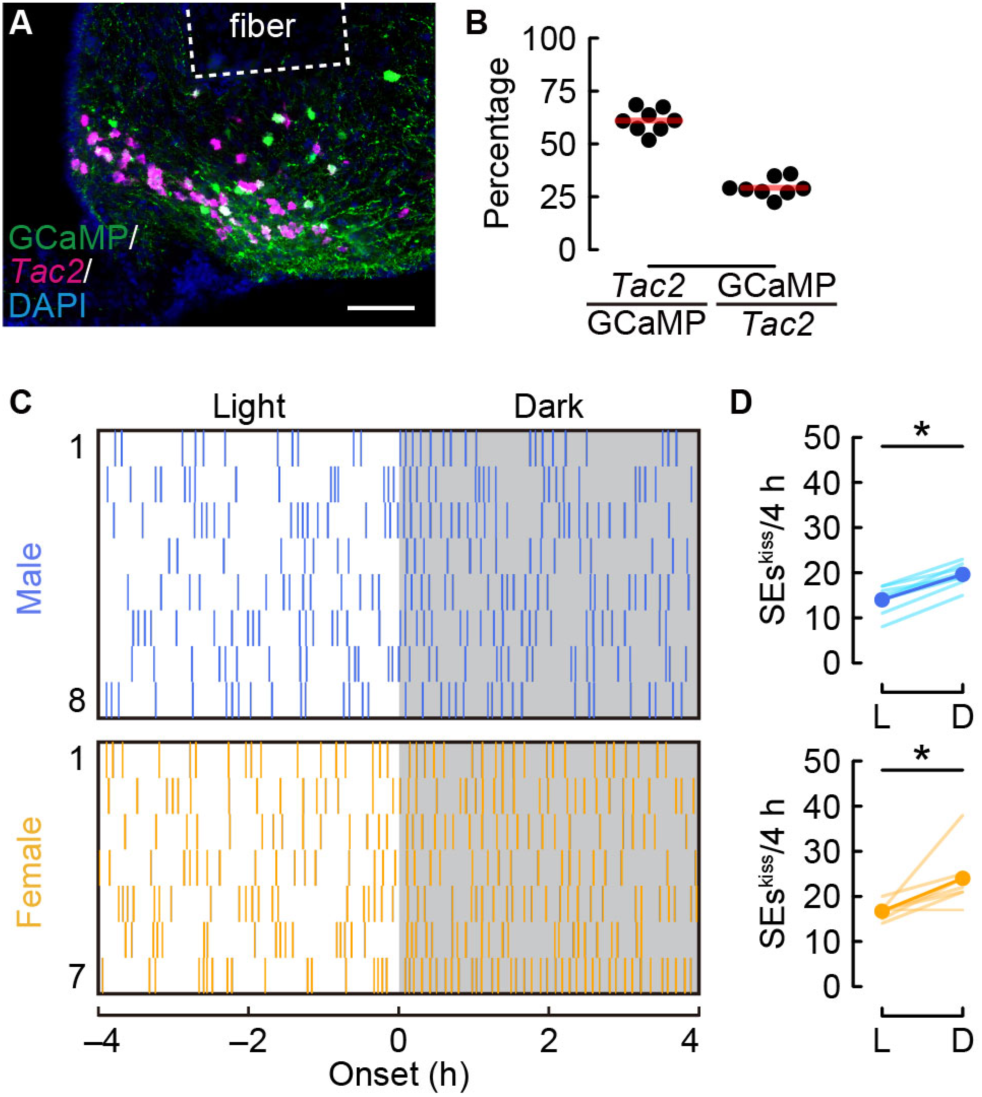
Additional analysis of SEs^kiss^ pattern in gonadectomized mice, related to Fig. 1. (A) Representative coronal sections of the ARC from *Kiss1-Cre* mice showing the optic fiber tract with *Tac2* mRNA (magenta; a marker of ARC^kiss^ neurons) and GCaMP6s (green; anti-GFP) counterstained with DAPI (blue). Scale bar, 100 μm. (B) Quantification of targeting specificity (*Tac2*+/GCaMP6s+) and efficiency (GCaMP6s+/*Tac2*+). *n* = 8. (C) Raster plots of SEs^kiss^ (vertical bars) in individual male (top) and female (bottom) mice during the last and first 4 h of the light and dark periods, respectively. Gray shading indicates the dark period. (D) Number of SEskiss during the last and first 4 h of the light (L) and dark (D) periods. \**p* < 0.05 using a two-sided paired *t*-test (D). *n* = 8 castrated males; *n* = 7 ovariectomized females (C, D).

**Fig. S2:**
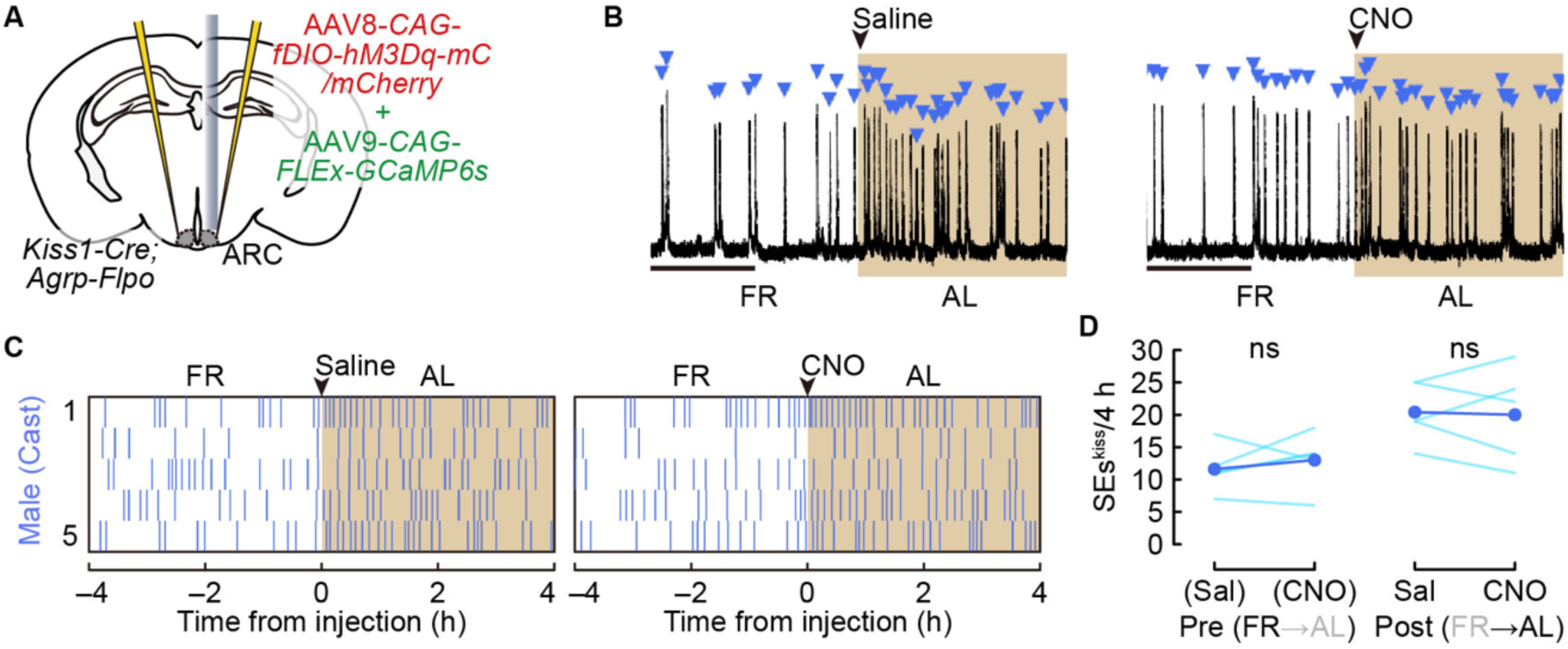
Chemogenetic activation of ARC^Agrp^ neurons does not affect SEs^kiss^ after FR removal in castrated adult male mice, related to Fig. 1. (A) Schematic of the experimental setup. (B) Representative photometry traces showing SEs^kiss^ (blue arrowheads) around FR removal. Arrowheads indicate the timing of intraperitoneal saline or CNO injection. Scale bars, 2 h. (C, D) Raster plots (vertical bars; C) and number (D) of SEs^kiss^ in individual male mice around FR removal, during the last and first 4 h of the FR and AL conditions, respectively. Brown boxes indicate the AL period after FR removal. No significant difference in SEs^kiss^ frequency was detected using a two-sided paired *t*-test (D). *n* = 5 castrated males.

**Fig. S3:**
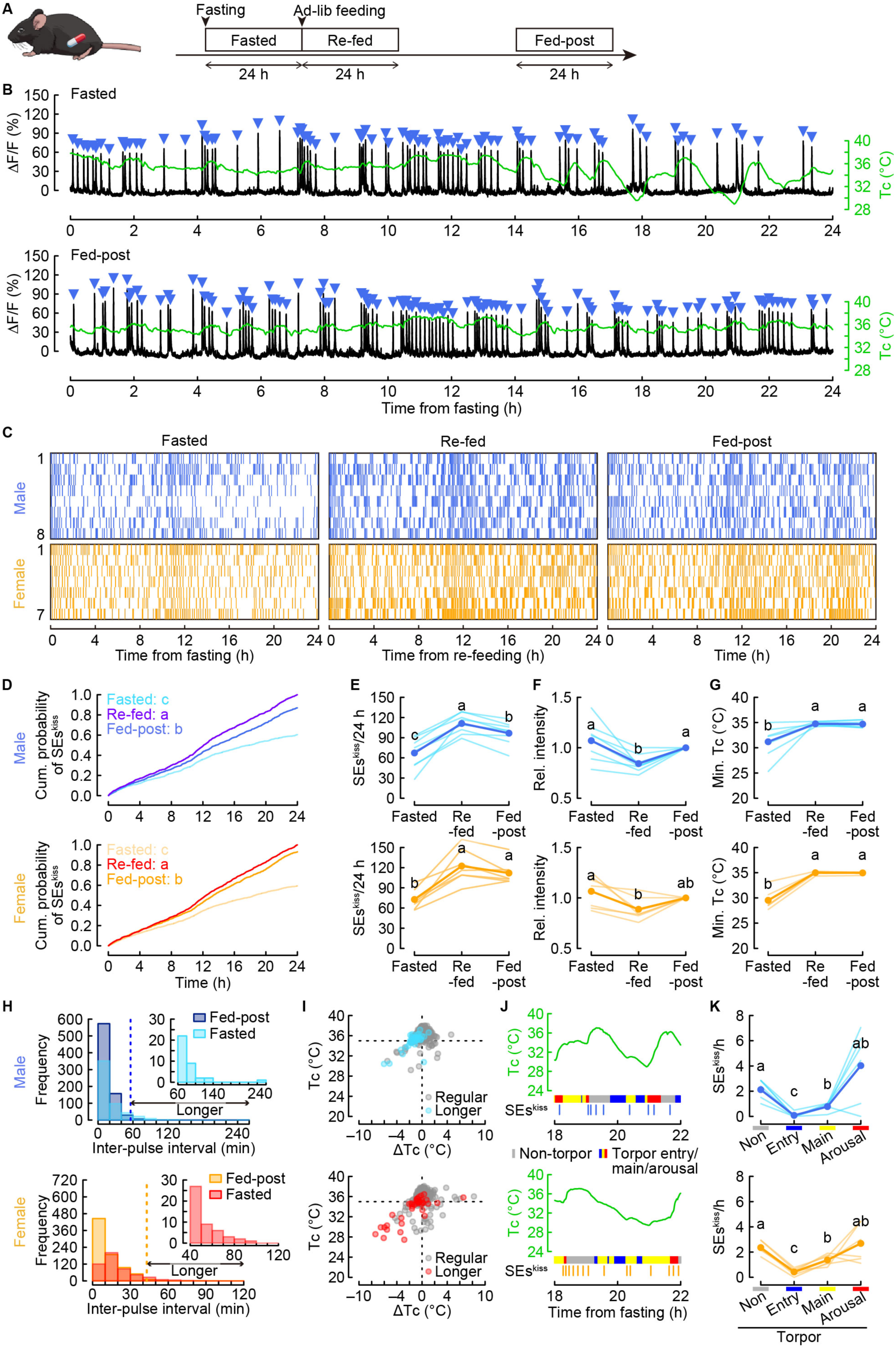
SEs^kiss^ are suppressed during daily torpor induced by 24-h fasting, related to Fig. 2. (A) Schematic of the experimental timeline. (B) Representative 24-h photometry traces (black) showing SEs^kiss^ (blue arrowheads) and Tc (green) under 24-h Fasted (top) and Re-fed (bottom) conditions in castrated male mice. (C) Raster plots of SEs^kiss^ (vertical bars) under Fasted, Re-fed, and Fed-post conditions. (D–F) Cumulative probability (D), number (E), and relative (rel) intensity, expressed as normalized ΔF/F (F), of SEs^kiss^. Different letters (a–c) indicate significant differences at *p* < 0.05 using the Kolmogorov–Smirnov test with Bonferroni correction (D). (G) Minimum Tc during photometry recordings (*n* = 8 castrated males; *n* = 7 ovariectomized females). Different letters (a, b) indicate significant differences at *p* < 0.05 using one-way repeated-measures ANOVA with Bonferroni-corrected post hoc tests (E–G). (H) Histogram of inter-pulse intervals under 24-h Fasted and Fed-post conditions. Dotted lines at 46 min (males) and 43 min (females) indicate the 99th percentile of inter-pulse intervals. Under 24-h fasting, intervals are categorized as regular or prolonged based on this threshold. (I) Scatter plots of ΔTc from the preceding SEs^kiss^ versus Tc at the current SEs^kiss^ in the Fasted group. Black and blue/red points denote regular and prolonged inter-pulse intervals, respectively. (J) Representative alignment of Tc (green), torpor phases (non-torpor, gray; entry, blue; maintenance, yellow; arousal, red), and SEs^kiss^ in the Fasted group. (K) Number of SEs^kiss^ across torpor phases in the fasting group. Different letters (a–c) indicate significant differences at *p* < 0.05 using one-way repeated-measures ANOVA with Bonferroni-corrected post hoc tests.

**Fig. S4:**
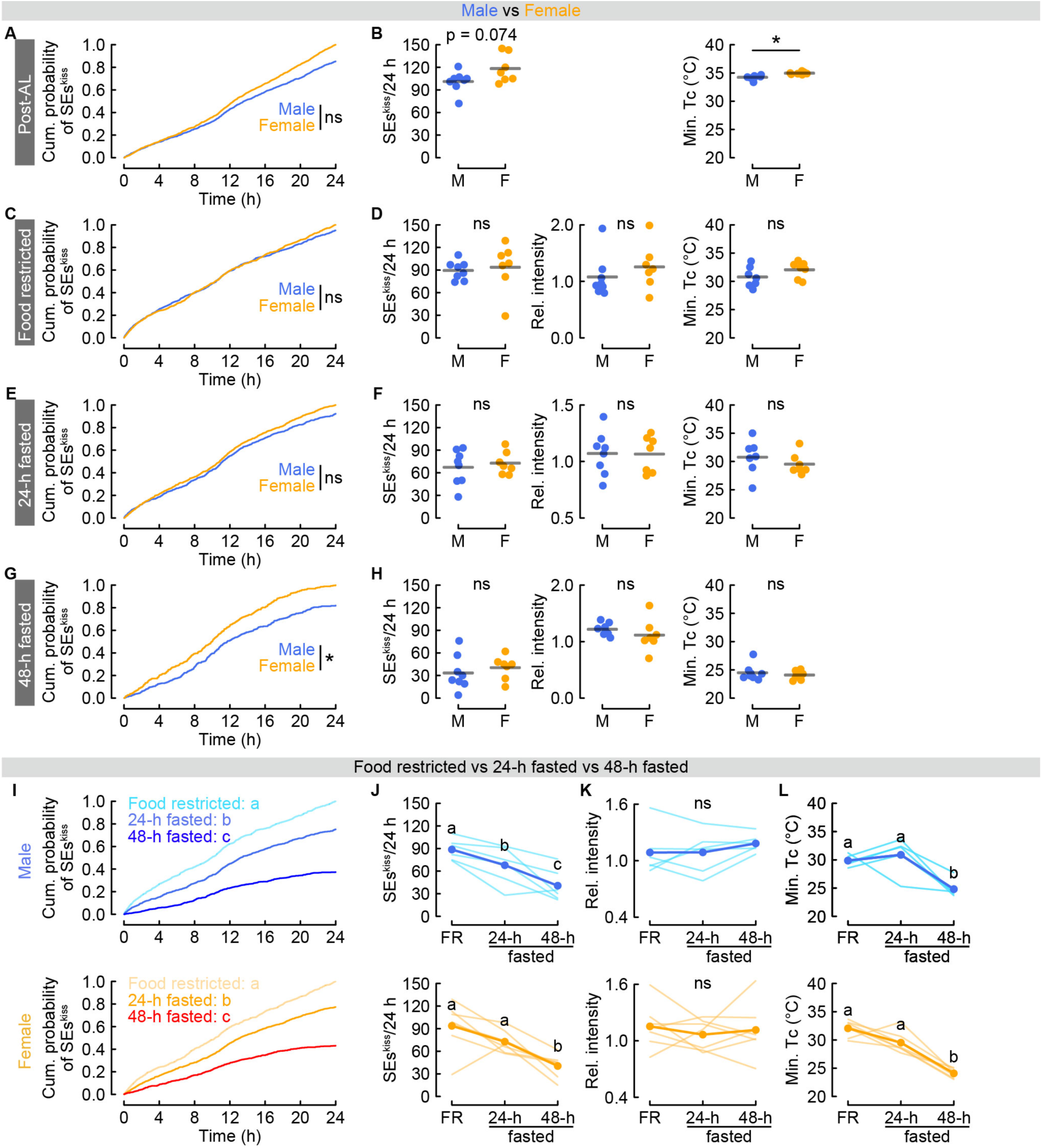
Comparison of different food-deprivation conditions, related to Fig. 2. (A, C, E, G) Cumulative probability distributions of SEs^kiss^ under post-AL (A), FR (C), 24-h fasted (E), and 48-h fasted (24–48 h window) (G) conditions in male and female mice. \**p* < 0.05 using the Kolmogorov–Smirnov test (G). (B, D, F, H) Number (left), relative intensity (normalized ΔF/F; middle) of SEs^kiss^, and minimum Tc (right) under post-AL (B), FR (D), 24-h fasted (F), and 48-h fasted (H) conditions in male and female mice. \**p* < 0.05 using a two-sided *t*-test (B). M, male; F, female. (I) Cumulative probability of SEs^kiss^ under FR, 24-h fasted, and 48-h fasted conditions. Different letters (a–c) indicate significant differences at *p* < 0.05 using the Kolmogorov–Smirnov test with Bonferroni correction. (J–L) Number (J), relative intensity (normalized ΔF/F) of SEs^kiss^ (K), and minimum Tc (L) in male (top) and female (bottom) mice under FR, 24-h fasted, and 48-h fasted conditions. Different letters (a–c) indicate significant differences at *p* < 0.05 using one-way repeated-measures ANOVA followed by Bonferroni-corrected post hoc tests. ns, not significant. *n* = 6 castrated males; *n* = 7 ovariectomized females.

**Fig. S5:**
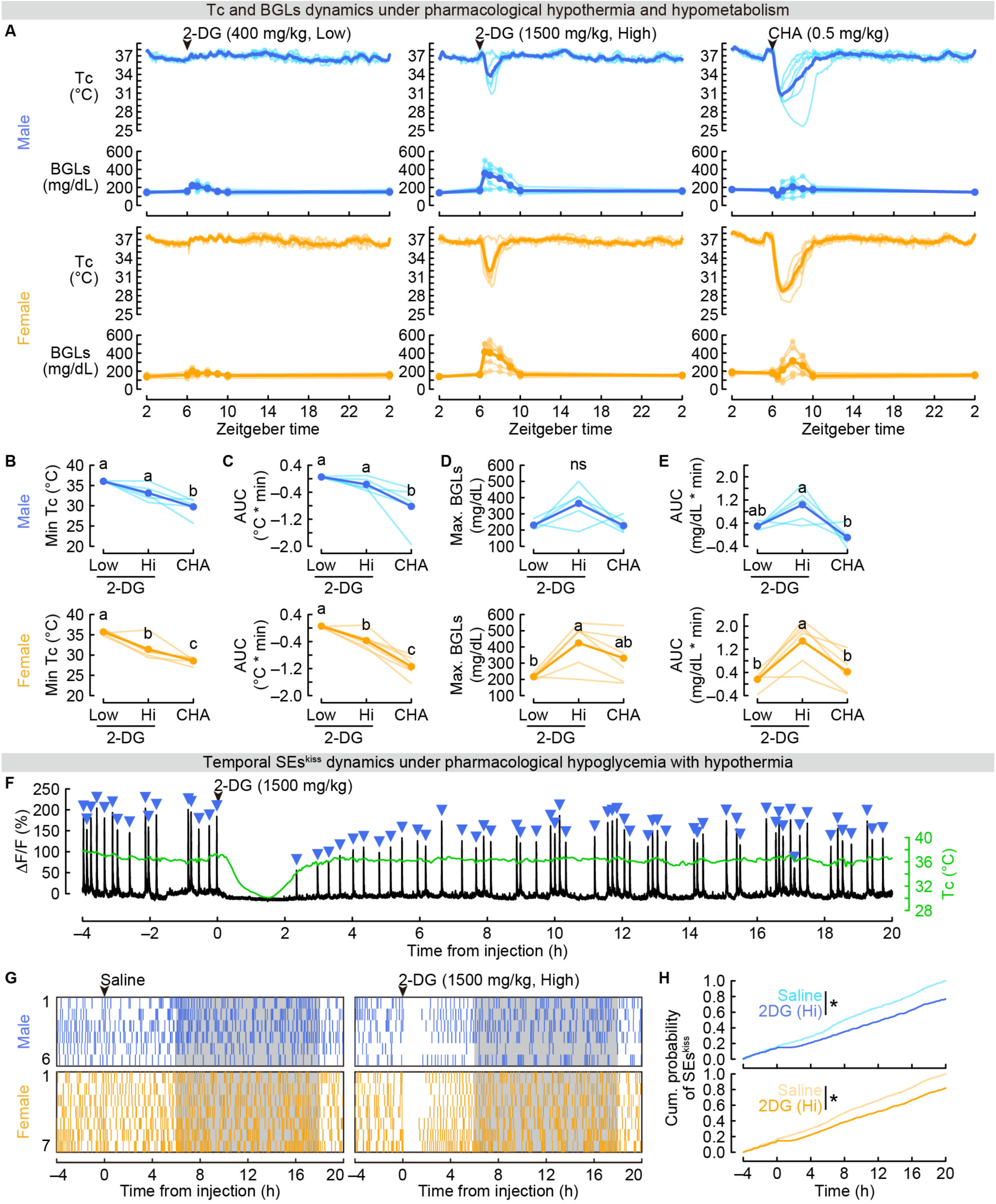
Pharmacologically induced hypothermia and BGLs in gonadectomized mice, related to Fig. 3. (A) Tc and BGLs following administration of low (left) or high (middle) doses of 2-DG and CHA (right). (B, C) Minimum Tc (B) and area under the curve (AUC) of Tc (C) during the 4-h period after drug administration. (D, E) Maximum BGLs (D) and AUC of BGLs (E) during the 4-h period after drug administration. Different letters (a–c) indicate significant differences at *p* < 0.05 using one-way repeated-measures ANOVA followed by Bonferroni-corrected post hoc tests. ns, not significant. *n* = 6 each for castrated males and ovariectomized females (A–E). (F) Representative 24-h photometric trace (black) showing SEs^kiss^ (blue arrowheads) and Tc (green) in a male mouse injected with 2-DG (1500 mg/kg, shown as black arrowheads). (G) Raster plots of SEs^kiss^ (vertical bars) in mice injected with saline (left) or 2-DG (right). (H) Cumulative probability of SEs^kiss^ in male (top) and female (bottom) mice injected with saline or 2-DG. \**p* < 0.05 using the Kolmogorov–Smirnov test with Bonferroni correction. *n* = 6 castrated males; *n* = 7 ovariectomized females (G, H).

**Fig. S6:**
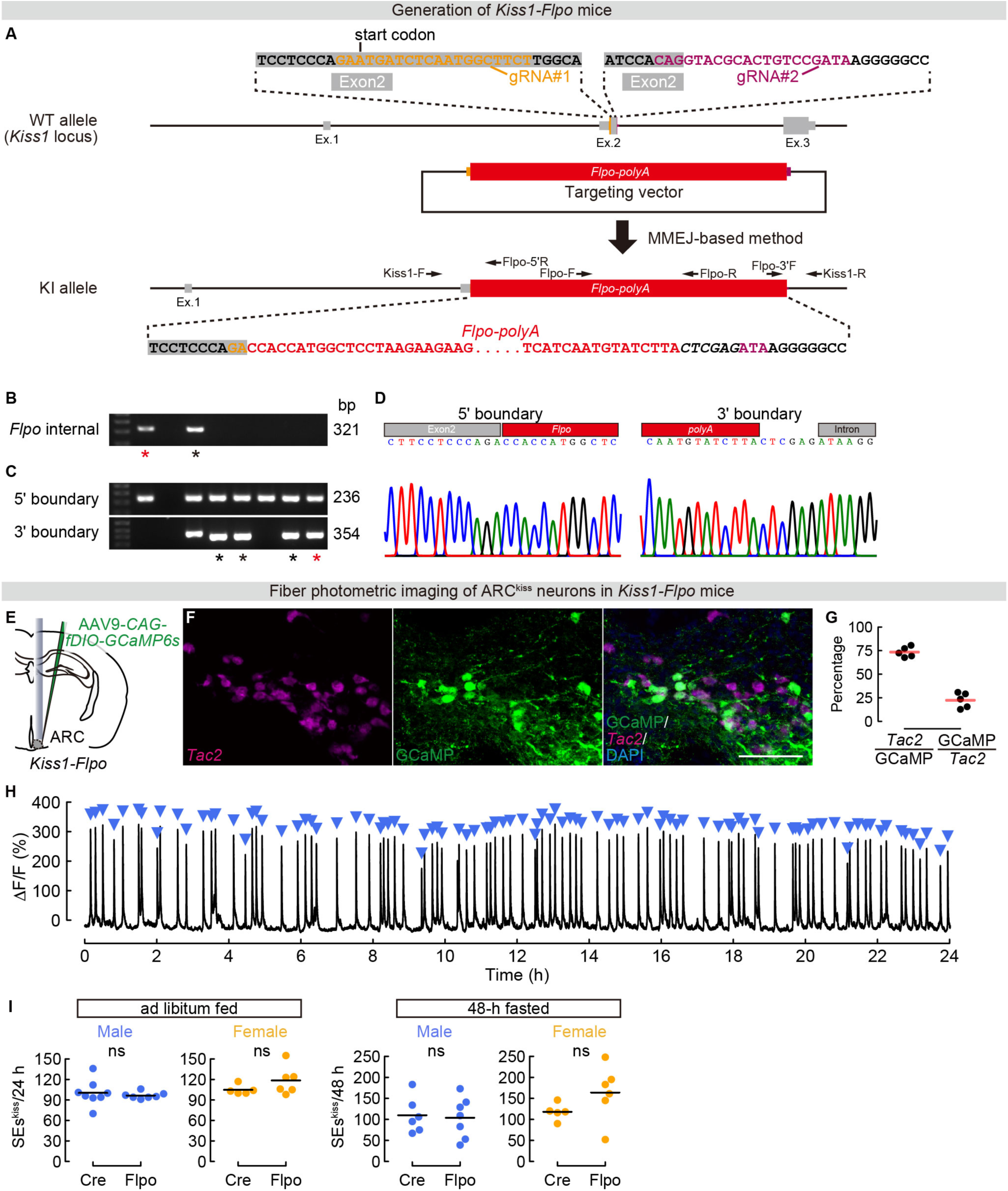
Generation of *Kiss1-Flpo* mice, related to Fig. 4. (A) Schematic of the knock-in strategy using the MMEJ-based method (*37*). The coding sequence of exon 2 at the *Kiss1* locus was replaced with an *Flpo-polyA* cassette. (B, C) Screening of *Kiss1-Flpo* founder mice. Genotyping PCR was performed using primers for the internal *Flpo* sequence (B) and the 5′ and 3′ junctions (C). Black asterisks indicate candidate F0 founders identified using PCR screening; red asterisks indicate established F0 lines confirmed by sequence analysis. PCR primers used were *Flpo-F*/*Flpo-R* (*Flpo* internal), *Kiss1-F*/*Flpo-5′R* (5′ junction), and *Flpo-3′F*/*Kiss1-R* (3′ junction). Expected PCR product sizes are indicated to the right of the gel images. See Methods for primer sequences. (D) Sequence confirmation of the 5′ and 3′ junction regions. (E) Schematic of the experimental setup. (F) Representative coronal brain sections of the ARC from *Kiss1-Flpo* mice showing *Tac2* mRNA expression (magenta; a marker of ARC^kiss^ neurons), and GCaMP6s (green; anti-GFP immunostaining), counterstained with DAPI (blue). Scale bar, 100 µm. (G) Quantification of specificity (*Tac2*+/GCaMP6s+) and efficiency (GCaMP6s+/*Tac2*+). *n* = 5 mice. (H) Representative 24-h photometric traces showing SEs^kiss^ (blue arrowheads) in castrated *Kiss1-Flpo* mice. (I) Number of SEs^kiss^ over 24 h under ad libitum feeding and 48-h fasting conditions in *Kiss1-Cre* and *Kiss1-Flpo* mice. ns, not significant using a two-sided non-paired *t*-test.

**Fig. S7:**
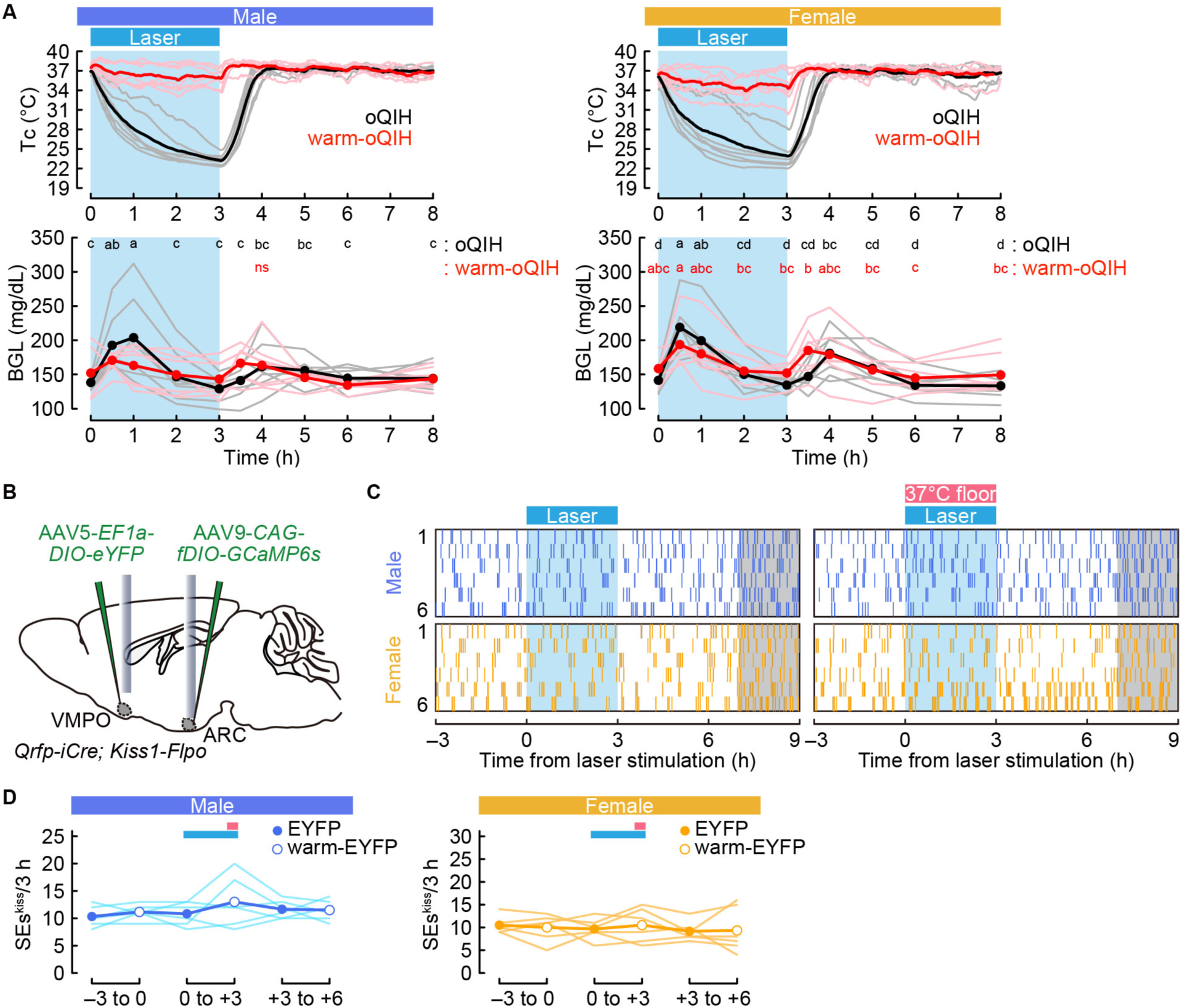
Tc and BGLs under oQIH and warm-oQIH conditions and EYFP control experiments, related to Fig. 4. (A) Tc and BGLs under oQIH and warm-oQIH conditions. Two-way repeated-measures ANOVA: QIH condition effect, ns; time effect, *p* < 0.01; interaction, *p* < 0.05 in males (left). QIH condition effect, *p* < 0.01; time effect, *p* < 0.01; interaction, *p* < 0.01 in females (right). Different letters (a– d) denote significant differences at *p* < 0.05 by one-way repeated-measures ANOVA followed by Bonferroni-corrected multiple comparisons within each group. ns, not significant. *n* = 7 each for castrated male and ovariectomized female mice. (B) Schematic of the control experiment expressing EYFP instead of ChR2 in Qrfp neurons. (C) Raster plots of SEs^kiss^ (vertical bars) in mice injected with control EYFP virus under laser illumination and warming-floor conditions. (D) Number of SEs^kiss^ per 3-h time window under these conditions. Two-way repeated-measures ANOVA: experimental condition effect, ns; time effect, ns; interaction, ns in both males and females. ns, not significant. *n* = 6 each for castrated male and ovariectomized female mice (B–D).

**Fig. S8:**
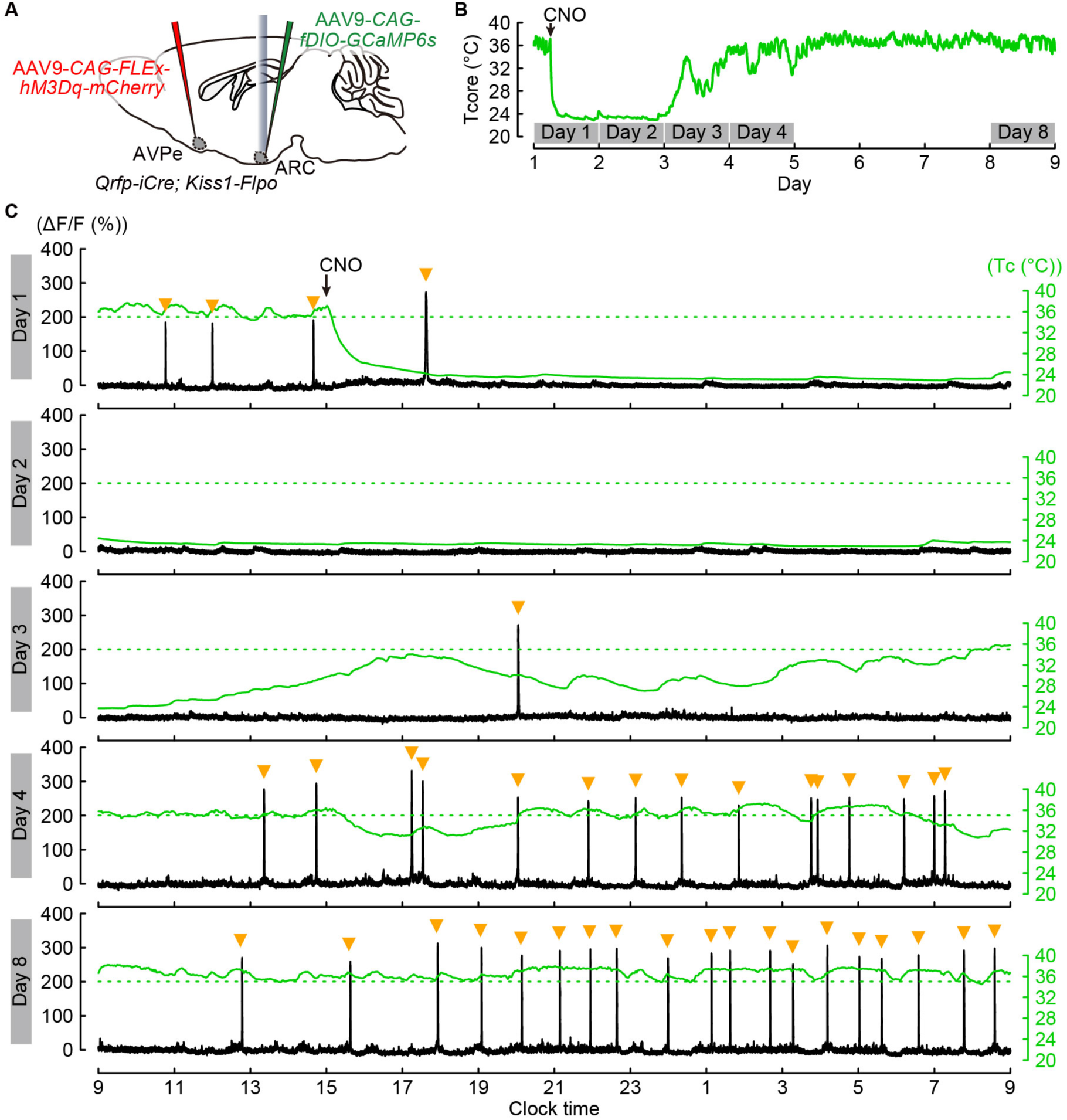
SEs^kiss^ are suppressed during chemogenetic QIH, related to Fig. 5. (A) Schematic of the experimental setup. (B) Representative Tc traces over 8 days. (C) Representative 24-h photometry traces (black) showing SEs^kiss^ (orange arrowheads) and Tc (green) under chemogenetic QIH in intact female mice. Arrows indicate the timing of CNO injection.

**Movie S1: Thermographic movies of mice during optogenetic QIH, related to Fig. 4**. Thermographic recording for 5 hours 14 minutes (12:32:26–17:46:26) was conducted using *Qrfp-iCre*; *Kiss1-Flpo* mice expressing hChR2(H134R) in the AVPe Qrfp neurons. The Qrfp neurons were stimulated at 2 Hz with a 10-ms pulse width for 3 hours (12:42:26–15:42:26).

**Movie S2: Thermographic movies of mice during warm-oQIH, related to Fig. 4**. Thermographic recording for 5 hours 14 minutes (12:32:24–17:46:24) was conducted using the same mice, as shown in Movie S1. The Qrfp neurons were stimulated at 2 Hz with a 10-ms pulse width for 3 hours (12:42:24–15:42:24). Simultaneously with the stimulation, the floor was heated to 37 °C.

